# Brain-wide organization of intrinsic timescales at single-neuron resolution

**DOI:** 10.1101/2025.08.30.673281

**Authors:** Yan-Liang Shi, Roxana Zeraati, International Brain Laboratory, Anna Levina, Tatiana A. Engel

**Author notes:** These authors contributed equally to this work (Yan-Liang Shi, Roxana Zeraati). Corresponding authors’.

## Abstract

Variations in intrinsic neural timescales across the mammalian forebrain reflect the anatomical structure and functional specialization of brain areas and individual neurons. Yet, the organization of timescales beyond the forebrain remains unexplored. We analyzed intrinsic timescales of single neurons across the entire mouse brain. Median timescales were up to fivefold longer in the midbrain and hindbrain than in the forebrain. Spatial patterns of gene expression predicted timescale variation at a resolution finer than brain-area boundaries. Across neurons, the diversity of timescales revealed a multiscale architecture, in which fast timescales determined regional differences in medians, while slow timescales universally followed a power-law distribution with an exponent near 2, indicating a shared dynamical regime across the brain consistent with the edge of instability or chaos. These organizing principles for the dynamics of single neurons across the brain provide a foundation for linking cellular activity with regional specialization and brain-wide computation.

---

The brain comprises hundreds of anatomically distinct regions, integrated into global functional networks that together support complex behavior across a broad range of timescales. Yet, the organizing principles of temporal information processing at the scale of the whole brain remain largely unknown. Anatomical structure provides the scaffold for neural dynamics, where regions distinguished by cell types, connectivity, and evolutionary origins form distinct functional modules. Determining the rules that organize dynamical properties of single neurons across the brain, which underpin the specialized functions of each region, is a critical step toward linking global brain structure to behavior.

In the mammalian cortex, intrinsic timescales of single neurons provide a distinctive signature that links neural dynamics to anatomical structure and behavioral function^1–3^. Intrinsic timescales quantify the temporal persistence of neural activity at rest, measured by the exponential decay rate of its autocorrelation^4^. Single-neuron timescales increase along the anatomical hierarchy of cortical areas^5–7^, defined by feedforward and feedback connectivity patterns ^8, 9^, and correlate with cortical gradients in pyramidal neuron spine density^10^ and the expression of synaptic receptor genes ^9,11^. Intrinsic timescales also align with the functional hierarchy of cortical processing: faster in early sensory areas that support rapid stimulus responses, and slower in association areas involved in evidence accumulation, working memory, and planning ^5–7,11–17^. Optogenetic inactivation of frontal areas disrupted evidence accumulation over longer timescales than inactivation of sensory areas, directly demonstrating the functional relevance of the timescale hierarchy^18^. Within individual frontal areas, neurons exhibit a broad diversity of timescales, with short- and long-timescale neurons encoding distinct cognitive variables during decision-making and working memory^19–23^. Thus, intrinsic timescales serve as a robust signature of cortical anatomical and functional hierarchy, suggesting they may provide a general marker for uncovering the organizing principles of temporal information processing across the whole brain.

However, beyond the cortical hierarchy, the brain-wide organization of single-neuron intrinsic timescales remains unexplored. Previous studies of single-neuron timescales have been confined to the forebrain, primarily the cortex and thalamus. In the thalamus, intrinsic timescales do not follow the hierarchical order implied by their cortical projections^7^. Nonetheless, thalamic timescales are generally faster than those in the cortex, consistent with the predominant direction of thalamocortical sensory input processing. This observation suggests that phylogenetically older subcortical areas, such as the midbrain and hindbrain, may also exhibit faster timescales than the cortex. Yet, these regions are critical for motor coordination and homeostatic regulation, functions that unfold over timescales of minutes to even hours^24–29^, and the cerebellum in particular may support slow, suprasecond processing^30^. Therefore, the logic of temporal processing at the brain-wide scale remains unclear and may be revealed through systematic measurements of single-neuron intrinsic timescales across the entire brain.

Using a dataset collected by the International Brain Laboratory (IBL)^31^, we analyzed intrinsic timescales of over ten thousand single neurons across 223 brain areas spanning the forebrain, midbrain, and hindbrain. We uncovered regional differences in timescales, their anatomical correlates, and their relationship to the encoding of cognitive variables. The diversity of timescales across neurons revealed a multiscale architecture, in which fast timescales drive regional differences, while slow timescales indicate a common dynamical regime across the brain. These principles governing single-neuron dynamics across the brain provide a foundation for establishing links between cellular activity, regional specialization, and brain-wide computation.

## Results

### Multiple intrinsic timescales in single-neuron activity

We analyzed intrinsic timescales of single-neuron activity in the IBL dataset of brain-wide Neuropixels recordings^31^. The recordings spanned 223 brain regions, covering all major structures of the mouse brain, including the isocortex (CTX), olfactory areas (OLF), hippocampal formation (HPF), cortical subplate (CTXsp), striatum (STR), pallidum (PAL), thalamus (TH), hypothalamus (HY), midbrain (MB), pons (P), medulla (MY), and cerebellum (CB). The brain-wide data were collected from 115 mice (80 males, 35 females) trained to perform the IBL decision-making task^32^ (Methods). During each recording session, mice first performed the task, followed by a recording of spontaneous activity. After applying the stringent IBL quality control metrics^31^, the dataset included spiking activity from 27,960 well-isolated single neurons, which we used in our analyses.

We measured intrinsic timescales from the autocorrelation of spontaneous spiking activity of single neurons. Spontaneous activity was recorded during a 10-minute period at the end of each session, while the mice were head-fixed and stationary in a holder, with no visual stimuli presented^32^. For each neuron, we computed the autocorrelation of spike counts in 5 ms time bins over the entire 10-minute recording (Methods).

Individual neurons exhibited diverse autocorrelation shapes both within and across brain areas (Fig. 1a). Timescales are typically estimated by fitting the autocorrelation with an exponential decay function, which appears as a straight line in logarithmic-linear coordinates, with a slope equal to the inverse of the timescale^5,12,19–22^. However, many neurons in our dataset displayed autocorrelations with multiple linear slopes, indicating that a single exponential decay was insufficient to capture their dynamics. These multiple slopes suggest the presence of several distinct timescales in the activity of individual neurons.

**Fig. 1.**
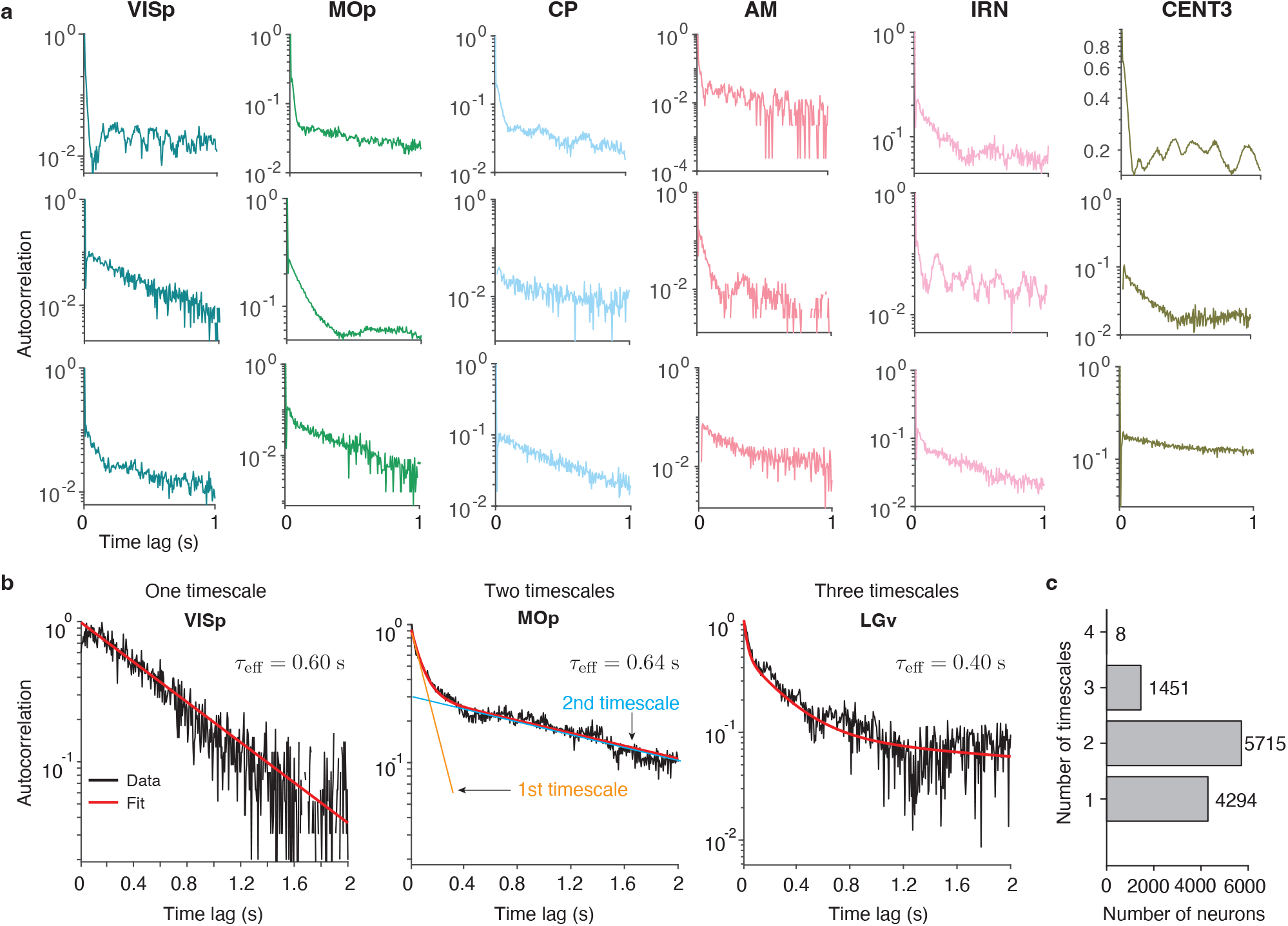
Multiple timescales in activity of single neurons across the mouse brain. **a**, Example single neurons exhibit diverse autocorrelation shapes both within brain areas (rows) and across areas (columns: VISp, primary visual area; MOp, primary motor area; CP, caudoputamen; AM, anteromedial nucleus; IRN, intermediate reticular nucleus; CENT3, lobule III). Different autocorrelation slopes in logarithmic-linear coordinates indicate distinct timescales. **b**, Autocorrelations of example single neurons best fit by one-timescale (left, VISp), two-timescale (center, MOp), and three-timescale (right, LGv: ventral part of the lateral geniculate complex) models (black: data, red: model fit). Multiple timescales in the model fit capture different linear slopes in the autocorrelation (center: orange and blue lines show exponential decays for the estimated first and second timescales, respectively). The effective timescale *τ*_eff_ summarizes the overall persistence of neural activity by matching the area under the multi-timescale autocorrelation curve to a single exponential decay with the effective timescale. **c**, Number of neurons best fit by models with one to four timescales. The two-timescale model provided the best fit for most neurons, while only eight neurons were best fit by the four-timescale model.

To capture multiple timescales in single-neuron dynamics, we fitted their autocorrelations with linear mixtures of exponential decay functions, one for each timescale (Methods). The contribution of each timescale to the autocorrelation was determined by its linear mixing coefficient. For each neuron, we fitted models with the number of timescales ranging from one to four. We then selected the optimal number of timescales using the Bayesian information criterion (BIC), with the constraint that each timescale contributes at least 1% to the overall autocorrelation shape (Methods). The selected model provided a good fit (coefficient of determination *R*^2^ *>* 0.5) for 11,468 out of 27,960 single neurons (41%), which we used in subsequent timescale analyses. This proportion of well-fitted neurons is consistent with previous studies of intrinsic timescales in single neurons^19, 22^. For the remaining neurons, the autocorrelation shape was dominated either by noise or oscillatory features that cannot be captured by exponential decay functions (Supplementary Fig. 1).

Individual neurons exhibited up to four distinct timescales in their spontaneous activity dynamics (Fig. 1b,c). The two-timescale model provided the best fit for the majority of neurons (*N* = 5, 715). The remaining neurons were predominantly best described by either one-timescale (*N* = 4, 294) or three-timescale models (*N* = 1, 451), with only eight neurons best fit by a four-timescale model. These results align with prior observations of multiple timescales in the activity of single neurons^6, 33^, and suggest that additional, slower timescales can be uncovered in neural dynamics when analyzing longer recording periods^34^.

### Variation of timescales across brain areas

The comprehensive coverage of brain areas in the IBL dataset enabled us to systematically examine how timescales vary across regions at the whole-brain scale. Neurons with one, two, or three timescales were distributed across all brain regions in similar proportions, with the exception of four-timescale neurons—most of which were found in the hindbrain (Supplementary Fig. 2). To facilitate comparison across neurons with varying numbers of timescales, we defined an effective timescale for each neuron as the weighted average of its individual fitted timescales, with weights given by their corresponding mixing coefficients (Methods). The effective timescale summarizes the multiple timescales contributing to a neuron’s dynamics by matching the area under the original multi-timescale autocorrelation curve to that of a single exponential decay with the effective timescale. Thus, the effective timescale captures the overall temporal persistence of neural activity, enabling direct comparison across neurons with varying numbers of fitted timescales.

**Fig. 2.**
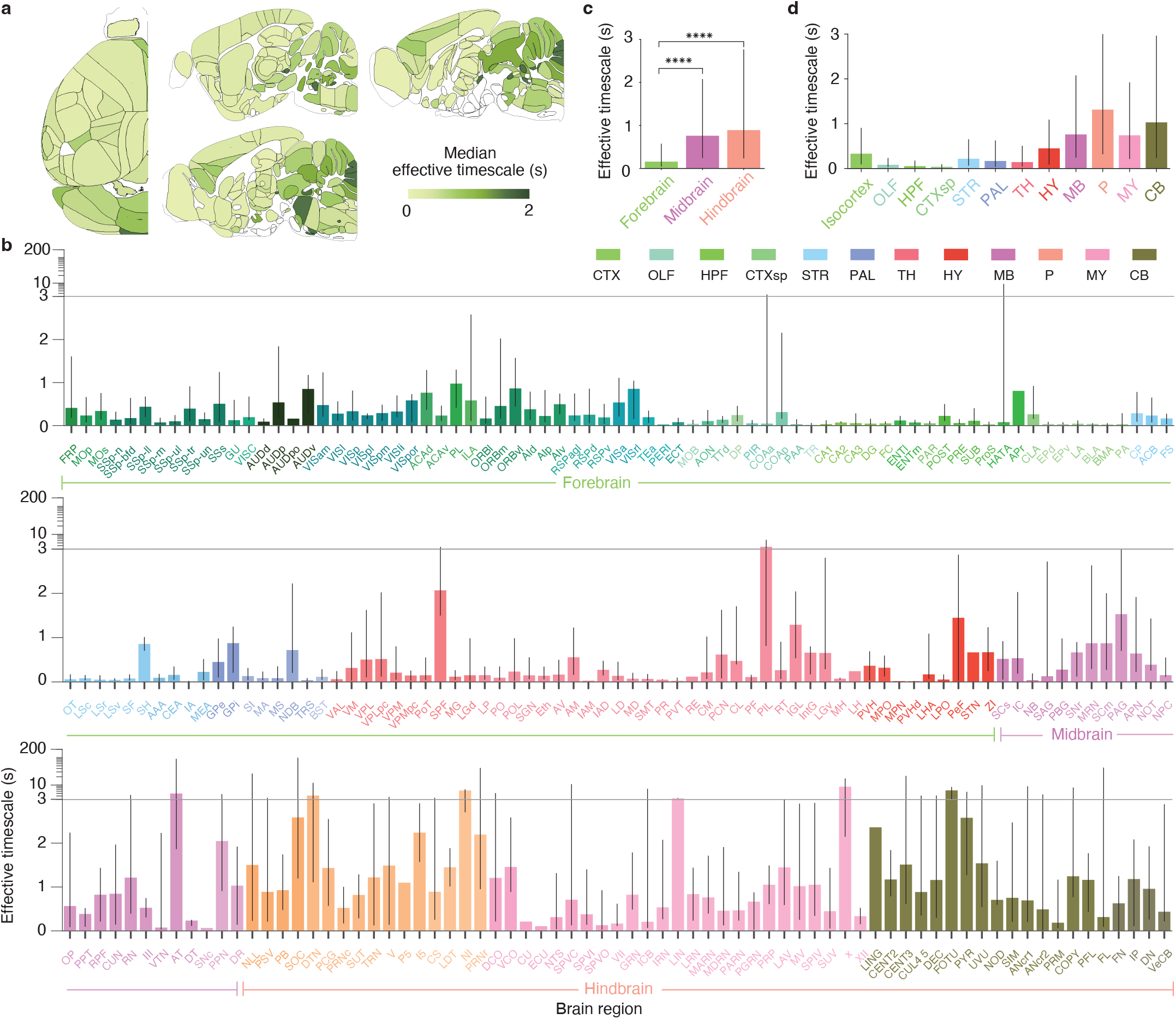
A map of intrinsic timescales across the mouse brain. **a**, Brain-slice plots (top and sagittal views) of the median effective timescale across brain regions. **b**, Effective timescales for all brain regions across the 12 major brain structures indicated by color (CTX, isocortex; OLF, olfactory areas; HPF, hippocampal formation; CTXsp, cortical subplate; STR, striatum; PAL, pallidum; TH, thalamus; HY, hypothalamus; MB, midbrain; P, pons; MY, medulla; CB, cerebellum). Bar height indicates the median, and error bars represent the 25th to 75th percentiles of the timescale distribution. The y-axis transitions from a linear to logarithmic scale (horizontal line) to accommodate the wide distribution of timescales. **c**, Effective timescales for the three major brain divisions: forebrain (including isocortex, olfactory areas, hippocampal formation, cortical subplate, striatum, pallidum, thalamus, and hypothalamus), midbrain, and hindbrain (including medulla, pons, and cerebellum). Bar height indicates the median (forebrain 0.16 s, *n* = 8, 399 neurons; midbrain 0.76 s, *n* = 914; hindbrain 0.89 s, *n* = 966), and error bars represent the 25th to 75th percentiles. Effective timescales were significantly longer in the midbrain and hindbrain than in the forebrain (two-sided Wilcoxon rank sum test, ^*∗∗∗∗*^*p <* 10^*−*10^). **d**, Effective timescales in 12 major brain structures. Bar height indicates the median, and error bars represent the 25th to 75th percentiles.

Median effective timescales varied widely across 223 brain areas spanning the forebrain, midbrain, and hindbrain (Fig. 2). Median effective timescales ranged from tens of milliseconds in some cortical, hippocampal, and striatal regions to several seconds in many midbrain, hindbrain, and cerebellar areas (Fig. 2a,b). Grouped by major brain division, neurons in the midbrain and hindbrain exhibited the longest median effective timescales—approximately five times longer than those in the forebrain (Fig. 2c, *p <* 10^*−*10^, two-sided Wilcoxon rank sum test; forebrain *n* = 8, 399; midbrain *n* = 914; hindbrain *n* = 966). This pattern was also evident when neurons were grouped into 12 major brain structures, with the shortest median effective timescales observed in olfactory areas, hippocampal formation, cortical subplate, and thalamus, and the longest timescales in the pons and cerebellum (Fig. 2d). We obtained similar results using the conventional method of fitting autocorrelations with a single exponential function (Supplementary Fig. 3), confirming the robustness of our findings to the choice of timescale estimation method. Our discovery of infra-slow intrinsic timescales in the midbrain and hindbrain suggests that these evolutionarily conserved brainstem structures may contribute to long-timescale integration of behavioral state information, rather than serving solely in the immediate control of movement.

**Fig. 3.**
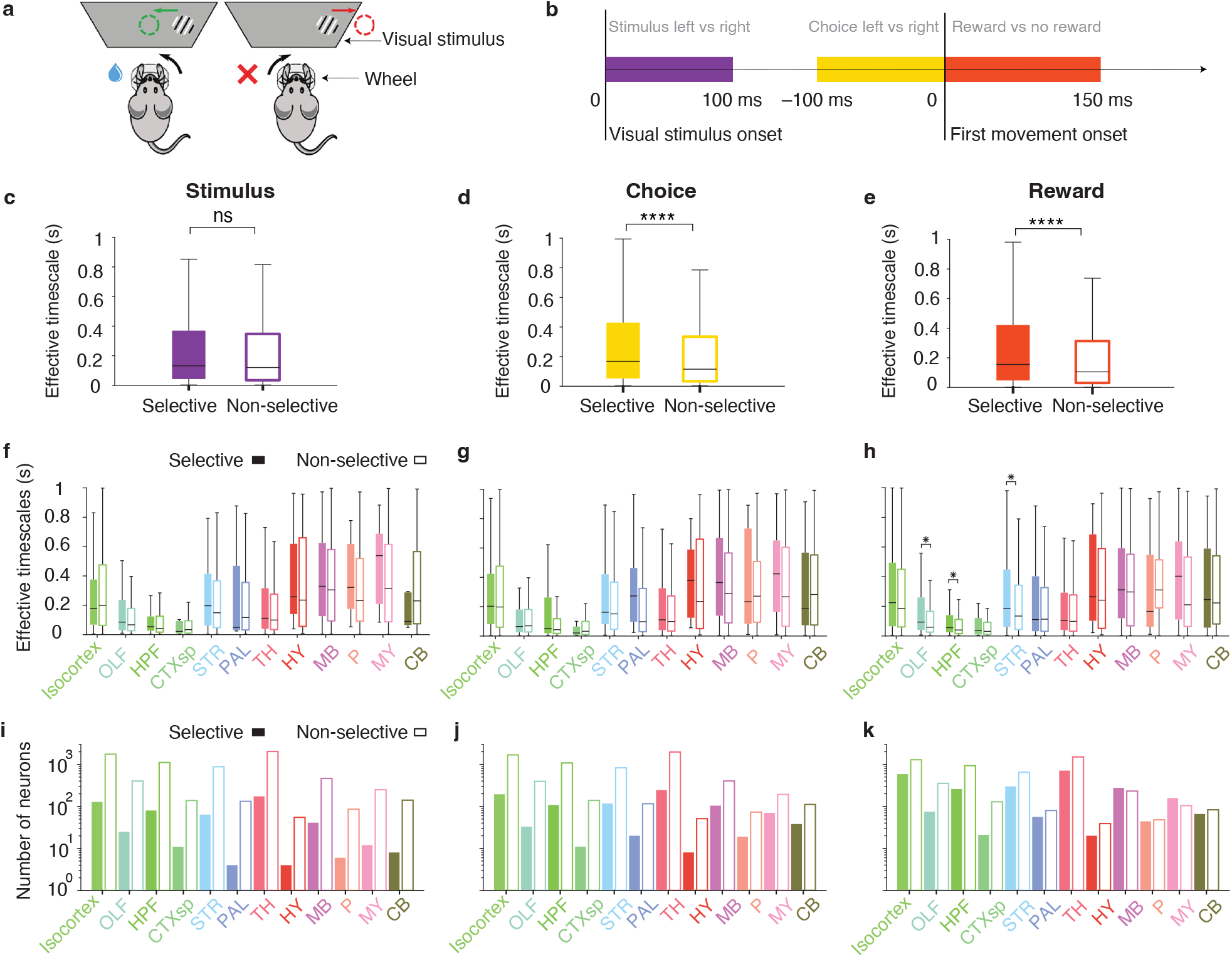
Relationship between single-neuron timescales and encoding of task variables. **a**, IBL decision-making task. Mice viewed a visual stimulus presented on either the left or right side of a screen. They received a reward for centering the stimulus by turning a wheel (left) and a timeout for incorrect responses (right). The illustration was adapted from ref.^31^. **b**, Time-windows used to quantify selectivity of single neurons to visual stimulus (purple, [0, 100] ms from stimulus onset), choice (yellow, [ − 100, 0] ms from movement onset), and reward (orange, [0, 150] ms from movement onset). **c**, Effective timescales did not differ significantly between neurons selective (filled bar) and non-selective (open bar) for the visual stimulus (*p* = 0.23, two-sided Wilcoxon rank sum test; *n* = 610 selective, *n* = 8, 196 non-selective). In box plots (c-h), center lines indicate medians; boxes span the 25th to 75th percentiles; whiskers extend to the nearest of 1.5× the interquartile range or the most extreme data point. **d**, Effective timescales were significantly longer in choice-selective neurons (filled bar) than in neurons non-selective for choice (open bar; *p <* 10^*−*10^, two-sided Wilcoxon rank sum test; *n* = 1, 067 selective, *n* = 7, 739 non-selective). **e**, Effective timescales were significantly longer in reward-selective neurons (filled bar) than in neurons non-selective for reward (open bar; *p <* 10^*−*10^, two-sided Wilcoxon rank sum test; *n* = 2, 847 selective, *n* = 5, 959 non-selective). **f**, Effective timescales did not differ significantly between neurons selective (filled bars) and non-selective (open bars) for the visual stimulus across all 12 major brain structures (*p >* 0.05, two-sided Wilcoxon rank sum test in each structure, Benjamini–Hochberg correction for multiple comparisons with FDR *α* = 0.01). **g**, Same as f for neurons selective and non-selective for choice. **h**, Effective timescales were significantly longer in reward-selective (filled bars) than in non-selective neurons (open bars) in olfactory areas, the hippocampal formation, and the striatum, with no significant differences in other brain structures (^*∗*^*p <* 0.05, two-sided Wilcoxon rank sum test in each structure, Benjamini–Hochberg correction for multiple comparisons with FDR *α* = 0.01). **i**, The number of neurons selective (filled bars) and non-selective (open bars) for the visual stimulus across 12 major brain structures. **j**, Same as i for choice. **k**, Same as i for reward.

### Relationship to task variable encoding

We further leveraged the brain-wide dataset to examine whether and how intrinsic timescales of individual neurons reflect their functional specialization in encoding behaviorally relevant variables. In association cortical areas, neurons with slower intrinsic timescales preferentially encode choice and working memory signals^19–23^. However, cognitive tasks recruit widespread neural populations well beyond association cortex, engaging subcortical structures that often exhibit some of the strongest and earliest encoding of task-related variables^35–37^. In particular, in the IBL decision-making task, visual stimulus representations first emerged in thalamic and cortical visual areas and gradually evolved into ramp-like stimulus activity in midbrain and hindbrain regions^31^. Choice- and reward-related signals were similarly widespread across the brain^31^. We therefore tested whether the selectivity of individual neurons for these task variables was related to their intrinsic timescales measured during spontaneous activity.

We quantified the selectivity of individual neurons to three task variables—stimulus, choice, and reward—in the IBL decision-making task. On each task trial, mice viewed a visual stimulus presented on either the left or right side of the screen and received a reward for centering the stimulus by turning a wheel (Fig. 3a, Methods). To assess neuronal selectivity, we used a condition-combined Mann-Whitney U-test, which compared spike counts across trials differing in one task variable (e.g., left vs. right stimulus side), within time windows aligned to the relevant behavioral event (Fig. 3b), while holding the other variables constant^31^. This approach allowed us to isolate the effect of each variable on neural activity, even when neurons were sensitive to multiple variables. We labeled neurons showing significant activity modulation for a given variable as selective (*p <* 0.05, condition-combined Mann-Whitney U-test; Methods), and all other neurons as non-selective.

For each task variable, we compared effective intrinsic timescales between selective and non-selective neurons. Intrinsic timescales were not significantly different between neurons selective versus non-selective for the visual stimulus (Fig. 3c; *p* = 0.23, two-sided Wilcoxon rank sum test; *n* = 610 selective, *n* = 8, 196 non-selective). By contrast, neurons selective for choice or reward exhibited significantly longer intrinsic timescales compared to their non-selective counterparts (Fig. 3d,e; *p <* 10^*−*10^, two-sided Wilcoxon rank sum test; choice: *n* = 1, 067 selective, *n* = 7, 739 non-selective; reward: *n* = 2, 847 selective, *n* = 5, 959 non-selective). These findings extend previous observations in association cortices^19–23^, demonstrating a relationship between intrinsic timescales and functional specialization of individual neurons throughout the brain.

To assess whether function-related differences in timescales were region-specific, we compared the effective timescales of selective and non-selective neurons separately within each of 12 major brain structures (Fig. 3f–k). Consistent with the whole-brain result, timescales did not differ significantly between neurons selective and non-selective for the visual stimulus in any structure (Fig. 3f, Supplementary Fig. 4, *p >* 0.05, two-sided Wilcoxon rank-sum test, Benjamini–Hochberg correction for multiple comparisons with FDR *α* = 0.01). While median effective timescales tended to be longer in choice-selective neurons than in non-selective ones across most structures, these differences were not statistically significant (Fig. 3g, Supplementary Fig. 4). In contrast, reward-selective neurons showed significantly longer timescales than non-selective neurons in olfactory areas (OLF), hippocampal formation (HPF), and striatum (STR) (Fig. 3h, Supplementary Fig. 4). This finding aligns with the known roles of these brain structures in reward processing: olfactory areas mediate sensory responses to water reward^38, 39^, the hippocampus contributes to reward-related memory^40^, and the striatum is involved in reward anticipation and outcome processing^41^. Overall, the preferential encoding of choice and reward by slow-timescale neurons may facilitate the long-term integration of these signals across trials to form prior expectations^42^.

### Anatomical correlates of timescales

Variation in intrinsic timescales may reflect anatomical differences in connectivity and cellular composition across the brain, motivating us to investigate this relationship using our brain-wide, cellular-resolution recordings. In the cortex, intrinsic timescales increase along the anatomical hierarchy defined by feedforward and feedback connectivity patterns, whereas thalamic timescales do not follow the hierarchical order implied by their cortical projections^6, 7^. Yet, the relationship between intrinsic timescales and anatomical features remains unexplored outside the forebrain—beyond the cortex and thalamus—or at finer spatial resolutions below the level of brain-area parcellations. We therefore tested the relationship between intrinsic timescales and anatomical features in two complementary ways: first, by confirming their association with connectivity-based hierarchical organization in the cortex and thalamus; and second, by correlating timescales with gene expression profiles across the whole brain at fine spatial resolution, reflecting continuous spatial variation in cellular and synaptic properties^10, 43^.

First, we evaluated the relationship between the timescales and anatomical hierarchy scores across cortical and thalamic areas^8^. Our dataset included 36 cortical and 22 thalamic areas (Supplementary Tables 1,2), providing near-complete coverage of the cortex and thalamus, unlike previous studies restricted to only a subset of regions (19 cortical and 7 thalamic)^6, 7^. Consistent with prior findings, median effective timescales were positively correlated with anatomical hierarchy scores in the cortex (Fig. 4a, Pearson correlation coefficient *r* = 0.41, *p* = 0.014, *n* = 36), but not in the thalamus (Fig. 4b, Pearson correlation coefficient *r* = 0.14, *p* = 0.526, *n* = 22). These results remained robust when restricting the analysis to the subset of areas studied previously^7^ (Supplementary Fig. 5). Moreover, when analyzed separately, most constituent timescales contributing to the effective timescale were positively correlated with anatomical hierarchy scores in the cortex, but not a single one exhibited a significant correlation in the thalamus (Supplementary Fig. 6). These results confirm that intrinsic timescales follow connectivity-defined hierarchical organization in the cortex, but not in the thalamus, based on a comprehensive dataset encompassing nearly all cortical and thalamic areas.

**Fig. 4.**
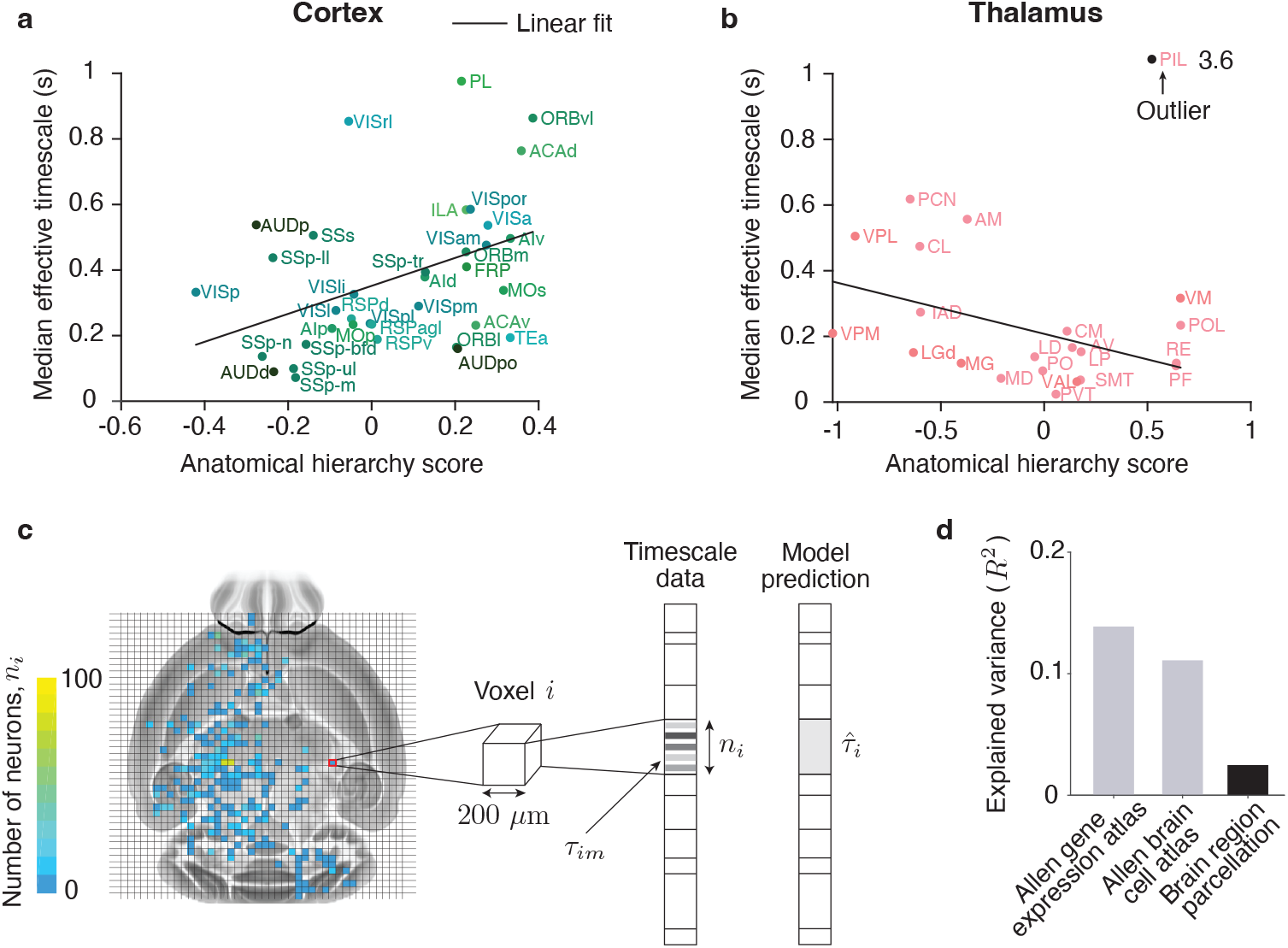
Relationship between intrinsic timescales and anatomical structure. **a**, Median effective timescales increased with the anatomical hierarchy scores in cortex (Pearson correlation coefficient, *r* = 0.41, *p* = 0.014, *n* = 36). Black line shows the linear fit. **b**, Median effective timescales did not significantly correlate with the anatomical hierarchy scores in thalamus (Pearson correlation coefficient, *r* = 0.14, *p* = 0.526, *n* = 22). One outlier with a large median effective timescale (PIL, 3.6 s) is clamped at the value 1 (black dot) for better visibility. Black line shows the linear fit with the outlier excluded. The correlation between median effective timescales and anatomical hierarchy scores was negative after excluding the outlier (*r* = −0.46, *p* = 0.034, *n* = 21). Abbreviations for all cortical and thalamic areas are listed in Supplementary Table 1. **c**, We used ridge regression to predict effective timescales of single neurons from the spatial densities of genes in the Allen Gene Expression Atlas^44^ or cell types in the Allen Brain Cell Atlas^43^, both of which cover the entire brain volume in cubic voxels with 200 *µ*m sides. Each voxel *i* contains *n*_*i*_ neurons with effective timescales *τ*_*im*_, where *m* = 1, … *n*_*i*_ (left). The regression predicts timescales from the anatomical features of the voxel where each neuron is located, so all neurons within a voxel share the same predicted value 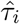 (right). **d**, Regression models based on anatomical features explained more variance in effective timescales than the baseline model, which predicted timescales solely from the brain region where each neuron was located (gene expression *R*^2^ = 0.14, cell-types *R*^2^ = 0.11, baseline *R*^2^ = 0.025).

Second, we tested whether anatomical features could predict intrinsic timescales across the entire brain at fine spatial resolution. To this end, we used two anatomical datasets: the Allen Gene Expression Atlas^44^, which provides spatial densities for 4,353 genes, and the Allen Brain Cell Atlas^43^, which provides spatial densities for 338 genetically defined cell types. Both datasets cover the three-dimensional brain volume in 3,955 cubic voxels with a side length of 200 *µ*m. We reduced the dimensionality of the gene expression data by projecting them onto the first 275 principal components (PCs), which explained 95% of variance. Thus, each voxel was characterized by a set of anatomical features: either 275 gene expression PCs or 338 cell-type densities. We then performed ridge regression across all neurons in the IBL data to predict their effective timescales based on the anatomical features of the voxel in which each neuron was located (Fig. 4c, Methods).

The regression models accounted for a substantial fraction of variance in the effective timescales across neurons (Fig. 4d, gene expression *R*^2^ = 0.14, cell-types *R*^2^ = 0.11, coefficient of determination, *n* = 11, 468). We confirmed that the prediction accuracy was not due to spurious correlations arising from independent spatial variation in both timescales and anatomical features^45^ (Supplementary Fig. 7, Methods). In comparison, prediction accuracy was considerably lower for the baseline regression model, which predicted effective timescales based solely on the brain region in which each neuron was located (*R*^2^ = 0.025, using 223 of 308 brain regions defined by the Allen Common Coordinate Framework^31, 46^; Methods). Since all models had a comparable number of predictors, these results indicate that intrinsic timescales vary spatially within brain-area boundaries at a resolution of 200 *µ*m, and that this fine-scale variation is partly explained by spatial differences in the composition of genetically defined cell types. In addition, different cell types contributed differentially to timescale prediction (Supplementary Fig. 8), offering potential insights into the molecular mechanisms that determine intrinsic timescales.

### Universal scaling of slow timescales

While median effective timescales varied fivefold across brain regions, we also observed extensive variability in timescales among individual neurons within each region (Fig. 2). Timescales of single neurons within each region spanned a broad range—from tens of milliseconds to several seconds—reflected by the wide error bars representing the interquartile range of their distribution (Fig. 2). Neurons with infra-slow timescales appeared even in regions with relatively fast median timescales, such as cortical areas. This pronounced variability and the presence of infra-slow timescales throughout all regions suggested that the timescale distributions might be heavy-tailed, leading us to analyze these distributions further.

We visualized the distributions of timescales across neurons within each major brain structure, as well as across neurons pooled from all regions (Fig. 5a). These distributions appeared as straight lines on logarithmic-logarithmic coordinates, suggesting power-law scaling of the timescales. To test for power-law scaling, we fitted the timescales with a power-law distribution and evaluated the hypothesis that the data follow a power law using a bootstrap-based statistical test^47^ (Methods). The results were consistent with a power-law distribution within each brain structure and in the data pooled across all regions (Supplementary Table 3). We further compared the goodness of fit of the power-law model against exponential and log-normal alternatives^48^ (Methods). Across all datasets, the power law provided a better fit than the exponential distribution and was statistically comparable to the log-normal distribution (Supplementary Table 5, Methods).

**Fig. 5.**
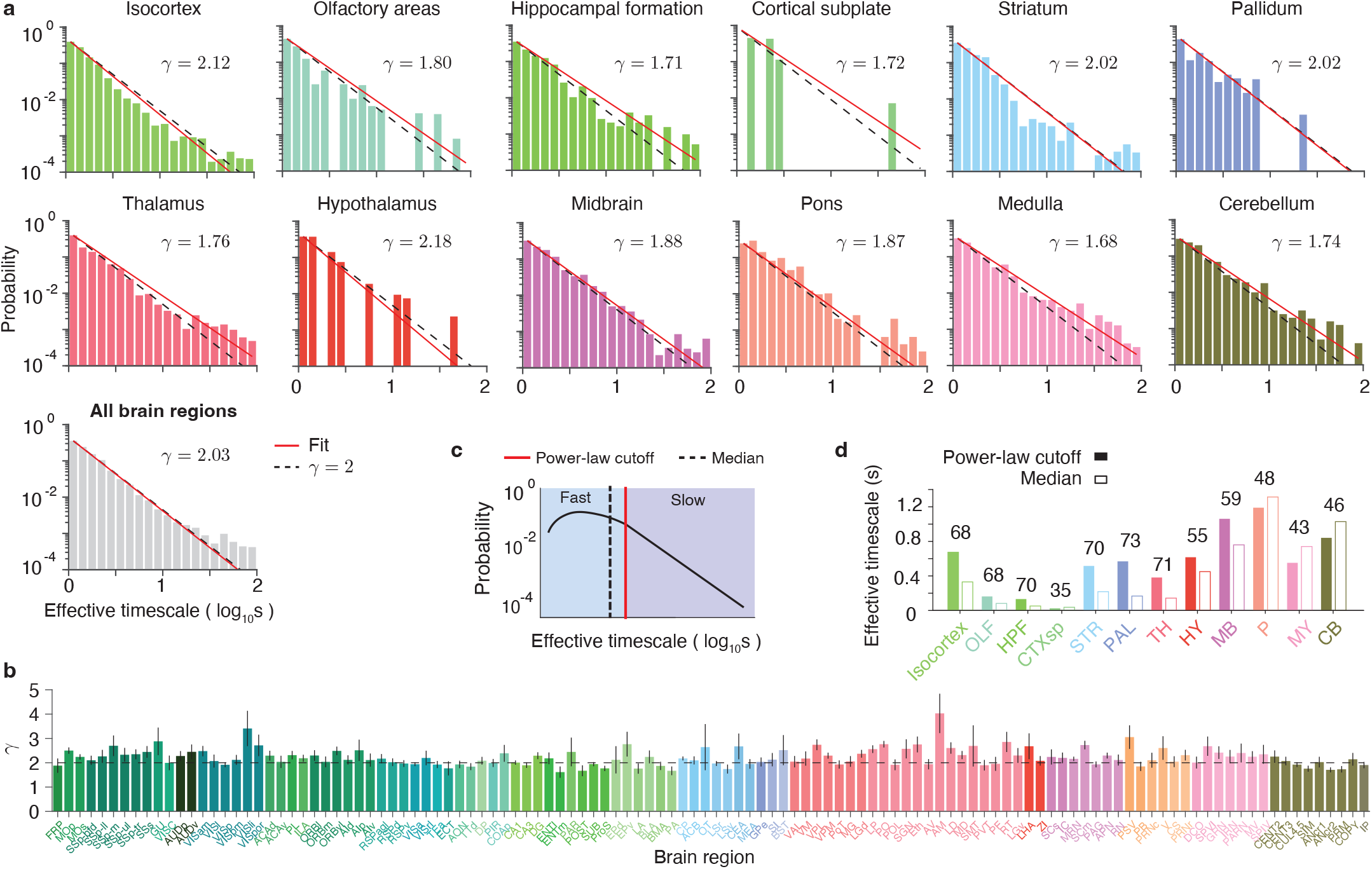
Universal scaling of timescales across the brain. **a**, The distributions of effective timescales across neurons within each major brain structure (colored) and across neurons pooled from all regions (gray) are well described by a power law with an exponent *γ* close to 2 (red line: best-fitting power law; dashed line: reference power law with exponent 2). **b**, The exponent *γ* of the best-fitting power-law for timescale distributions was near 2 in most brain regions, which had sufficient data for fitting (more than 40 neurons). Error bars represent 95% confidence intervals. **c**, The power-law distribution is defined only above a cutoff value determined during the fitting procedure. The cutoff divides timescales into fast (below the cutoff, blue) and slow (above the cutoff, purple), with only the slow timescales following the power-law. **d**, In all brain structures, cutoff values (solid bars) either exceeded the median timescale (open bar) or fell just below it. Numbers above bars indicate cutoff percentiles.

The scaling exponent *γ* of the power-law fit was close to 2 in all brain structures and in the data pooled from all regions (Fig. 5a), indicating a universal scaling of timescales throughout the brain. To further test this universality, we fitted timescales separately for neurons in each brain area with sufficient data (more than 40 neurons) and found that the scaling exponent remained close to 2 across most areas (Fig. 5b). The scaling exponent also remained near 2 when fitting timescales separately for all neurons recorded in each session (Supplementary Fig. 9a), indicating that the observed scaling was not an artifact of differences across sessions, animals, or laboratories. The observed scaling was also robust to the timescale estimation method: timescales estimated using a single-exponential fit were likewise best described by a power-law distribution with an exponent near 2 in all major brain structures and in the data pooled from all regions (Supplementary Fig. 9b, Supplementary Table 4). In addition, timescales of individual neurons were only weakly correlated with their firing rate (Pearson correlation coefficient, *r* = 0.04, *p* = 1 · 10^*−*4^, *n* = 11, 468), suggesting that the power-law scaling was not a trivial consequence of a heavy-tailed firing rate distribution^49^. Together, these analyses reveal a robust and widespread power-law scaling of intrinsic timescales with a consistent exponent near 2 throughout the brain.

Despite this universal scaling, median timescales varied fivefold across brain regions, raising the question of how such variability can be reconciled with a common power-law structure. The power-law distribution is defined only above a minimum value—the cutoff—which is determined during the fitting procedure^48^ (Methods). This cutoff divides timescales into fast (below the cutoff) and slow (above the cutoff), with only the slow timescales following the power-law (Fig. 5c, Supplementary Fig. 10). In all brain structures, the power-law cutoff either exceeded the median timescale or fell just below it, falling above the 43rd percentile in 11 of the 12 brain structures (Fig. 5d). Thus, variations in median timescales across brain regions were primarily driven by differences in the fast timescales below the power-law onset (Fig. 5c), while the slow timescales consistently followed the universal power-law scaling.

These results reveal a multi-scale architecture of intrinsic timescales throughout the brain, with fast timescales driving regional differences and slow timescales exhibiting universal scaling. Neurons with fast and slow timescales showed similar relationships to the encoding of behavioral variables: neurons selective for choice or reward exhibited longer timescales than non-selective neurons, whereas timescales did not differ between neurons selective and non-selective for visual stimulus (Supplementary Fig. 11). Regional differences in fast timescales may reflect variations in cellular properties across brain areas, as suggested by correlations between median timescales and anatomical features, while the universal scaling of slow timescales may indicate a common regime of network dynamics throughout the brain.

### Mechanisms for multiscale architecture

We discovered a multi-scale architecture of intrinsic timescales across the mouse brain, where fast timescales vary widely and correlate with anatomical features across regions, while slow timescales exhibit universal power-law scaling. The variation and universality in fast and slow timescales suggest possible underlying network mechanisms, which we explored using recurrent network models.

We considered two types of recurrent networks: linear^50, 51^ and nonlinear^52^. In both linear and nonlinear networks, we investigated which connectivity structures and dynamical regimes can give rise to universal power-law scaling of slow timescales, and which parameters control the power-law cutoff, and thereby the fast and median timescales.

In linear networks, a power-law distribution of timescales can arise from dynamics operating near the edge of instability^51^. To maintain stable dynamics—fluctuations around a fixed point driven by input noise— the eigenvalue spectrum of the connectivity matrix must remain bounded by zero, the threshold of instability (Fig. 6a). This condition can be satisfied when the connectivity is not heavy-tailed, resulting in a finite spectral range. In this case, timescales are given by the inverse of the eigenvalues, and the timescale distribution *g*(*τ*) is analytically related to the eigenvalue spectrum *f* (*λ*) by *g*(*τ*) = *f* (1*/τ*)*/τ*^2^ (Methods)^51^. When the synaptic gain *g* is tuned to bring the spectrum to the edge of instability, where the smallest eigenvalue approaches zero, extremely long timescales emerge. Approximating *f* (*λ*) near zero by the leading term of its power-series expansion, *f* (*λ*) ∝ *λ*^*ζ*^ as *λ* → 0 (with *ζ >* −1, Fig. 6a), we find that the slow timescales follow a power-law distribution *g*(*τ*) ∝ 1*/τ*^2+*ζ*^ with the exponent 2 + *ζ*. The inverse of the largest eigenvalue for which this approximation still holds defines the cutoff of the power-law distribution of timescales.

**Fig. 6.**
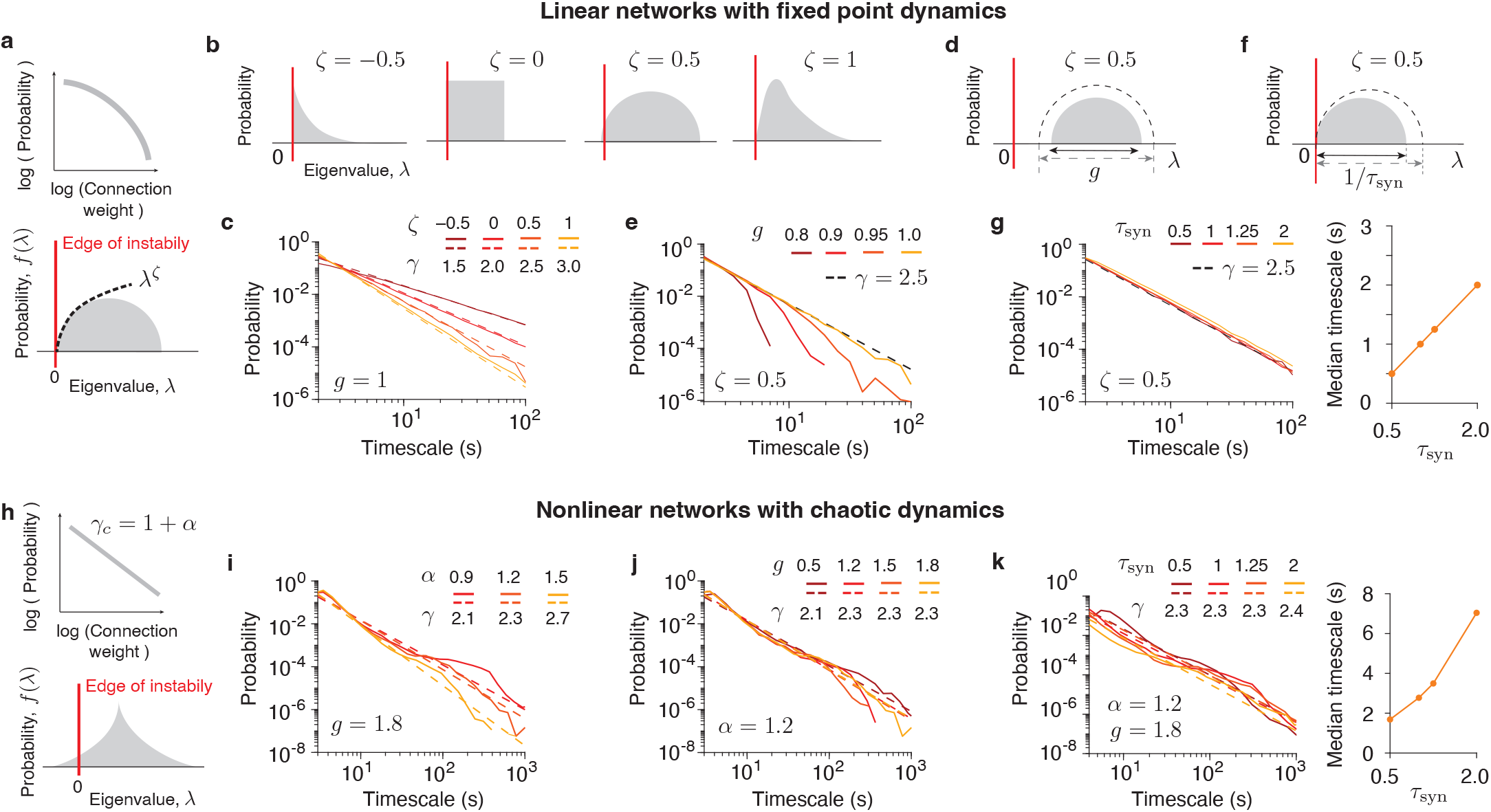
Network mechanisms for multiscale architecture. **a**, In linear networks, connection weights have a non-heavy-tailed distribution (upper panel). The connectivity eigenvalue distribution is bounded and approximated near zero by the leading term of its power-series expansion: *f* (*λ*) ∝ *λ*^*ζ*^ as *λ* → 0 (lower panel). **b**, Example connectivity eigenvalue distributions characterized by distinct values of *ζ* near zero: *ζ* = −0.5, gamma distribution with a shape parameter 0.5; *ζ* = 0, uniform distribution; *ζ* = 0.5, semi-circle distribution; *ζ* = 1, gamma distribution with a shape parameter 2. **c**, At the edge of instability, linear networks generate a power-law distribution of timescales with exponent *γ* = 2 + *ζ* (solid lines: simulations of network timescales computed as the inverses of connectivity eigenvalues sampled from distributions in panel b; dashed lines: power law with exponent derived from theory). **d**, The synaptic gain *g* controls the distance to the instability threshold by scaling the range of the connectivity eigenvalue distribution. **e**, Reducing *g* shifts the network further from the instability threshold, causing power-law collapse (solid lines: simulations of network timescales for *ζ* = 0.5; dashed line: reference power law with *γ* = *ζ* + 2). **f**, The synaptic time constant *τ*_syn_ scales the range of the connectivity eigenvalue distribution without affecting the distance to the instability threshold. **g**, Varying *τ*_syn_ does not affect the power-law exponent, while modulating the cutoff and thereby median timescale (solid lines: simulations of network timescales for *ζ* = 0.5; dashed line: reference power law with *γ* = *ζ* + 2). **h**, In nonlinear networks, the heavy-tailed connectivity with *α*-stable distribution has a power-law tail with exponent *γ*_*c*_ = 1 + *α* (upper panel). The connectivity eigenvalue spectrum extends beyond the linear instability threshold (lower panel). **i**, In the chaotic regime, nonlinear networks generate a power-law distribution of timescales with exponent *γ* matching that of the connectivity distribution (solid lines: network simulations for varying connectivity scaling parameter *α*; dashed lines: best-fitting power law with exponent *γ*; *g* = 1.8). **j**, The scaling exponent of the timescale distribution is independent of the synaptic gain *g* (solid lines: network simulations for varying *g*; dashed lines: best-fitting power law with exponent *γ*; *α* = 1.2). **k**, Varying *τ*_syn_ does not affect the power-law exponent, while modulating the median timescale (solid lines: network simulations for varying *τ*_syn_; dashed lines: best-fitting power law with exponent *γ*; *α* = 1.2, *g* = 1.8).

We confirmed our theory through simulations of linear networks with various connectivity eigenvalue distributions that exhibited different power-series behavior near zero, characterized by distinct values of *ζ* (Fig. 6b). From each distribution, we sampled *n* = 1, 000 eigenvalues and computed their inverses as network timescales, which closely approximate the distribution of single-neuron timescales in linear networks (Supplementary Fig. 12, Methods). As predicted by theory, setting the networks at the edge of instability produced a power-law distribution of timescales with exponent *γ* = 2 + *ζ* (Fig. 6c). The synaptic gain *g* controls the proximity to the instability threshold, and must be finely tuned near instability to observe the power law (Fig. 6d,e). The power-law cutoff can be modulated by varying the synaptic time constant parameter *τ*_syn_, which scales the range of the eigenvalue distribution while keeping the network at the edge of instability (Fig. 6f, Methods). Varying *τ*_syn_ does not affect the power-law exponent, while modulating both the cutoff and median timescales by a factor of *τ*_syn_ (Fig. 6g, Methods).

In nonlinear networks, a power-law distribution of timescales can emerge from heavy-tailed connectivity in the chaotic regime. Nonlinear networks can sustain chaotic dynamics when the eigenvalue spectrum of the connectivity extends beyond the linear instability threshold^52^ (Fig. 6h). When the synaptic gain *g* exceeds a critical value, the network transitions into a chaotic regime that generates a heavy-tailed distribution of timescales^52^.

To investigate the mechanisms governing the timescale distribution, we simulated nonlinear networks with heavy-tailed connectivity. We sampled connectivity weights from an *α*-stable distribution (0 *< α <* 2), which exhibits an asymptotic power-law tail with exponent *γ*_*c*_ = 1 + *α*. Simulations revealed that the timescale distribution also followed a power law with scaling exponent *γ* = 1 + *α*, matching that of the connectivity distribution (Fig. 6i). The scaling exponent of the timescale distribution was independent of the synaptic gain *g* (Fig. 6j), providing a mechanism to generate a universal power-law distribution of timescales without tuning *g*. As a control, networks with Gaussian connectivity^53^ (corresponding to an *α*-stable distribution with *α* = 2) exhibited timescale distributions that were not heavy-tailed (Supplementary Fig. 13), reinforcing the link between power-law scaling of connectivity and timescales. Similar to linear networks, the synaptic time constant *τ*_syn_ controlled the power-law cutoff and the median timescale without altering the scaling exponent (Fig. 6k).

These two models offer alternative mechanisms for the universal scaling of timescales: in linear networks, it emerges from non-heavy-tailed connectivity at the edge of instability, whereas in nonlinear networks, it arises from heavy-tailed connectivity in the chaotic regime. Heavy-tailed connectivity has been observed in several biological neural networks^54–56^, suggesting that a common connectivity scaling across areas may underlie the universal scaling of timescales in the mouse brain. In both models, the synaptic time constant sets the powerlaw cutoff and median timescale without altering the power-law exponent. This mechanism explains how universal scaling can persist despite variability in cellular and synaptic composition across areas and aligns with our finding that median timescales correlate with anatomical features. Whole-brain, cellular-resolution connectomes^57^ will enable direct assessment of the scaling properties of connectivity distributions, thereby constraining the network mechanisms for universal scaling of timescales.

## Discussion

We uncovered a brain-wide organization of intrinsic timescales, with the midbrain and hindbrain expressing up to fivefold longer timescales than the forebrain. The distribution of timescales across neurons revealed a multiscale architecture, in which fast timescales drive regional differences, while slow timescales follow a universal power law.

The infra-slow timescales in the midbrain and hindbrain may reflect the intrinsic capacity of these evolutionarily conserved brainstem structures to integrate information over extended periods. Indeed, both regions contribute to the integration of sensory and motor inputs across long timescales^26, 27^ and are critically involved in motor control, including speed regulation and gait selection^28^. The hindbrain additionally regulates bodily homeostasis, such as breathing rhythms and blood circulation^24^, as well as nutrient intake and gastrointestinal satiation signals^29^. These functions require integration and coordination over timescales ranging from seconds to minutes or even hours, consistent with the prevalence of neurons with infra-slow intrinsic dynamics. As brainstem areas are reciprocally connected with the rest of the brain^25^, they may modulate neural activity globally, broadcasting internal states and physiological needs to influence information processing throughout the brain^58, 59^.

Our finding that neurons selective for choice and reward exhibit longer timescales than non-selective neurons supports the view that intrinsic timescales reflect a neuron’s engagement in cognitive computations. This association, which we identified in brain-wide mouse data, aligns with similar reports from primate frontal cortex^19–23^, although it may depend on the timescale estimation method^13^. The causal link between intrinsic timescales and functional specialization of single neurons can be directly tested in recurrent network models trained to perform cognitive tasks^60–64^. These models provide a powerful framework for generating testable hypotheses for future experiments probing the causal role of single-neuron timescales in computation.

Across the nearly complete set of cortical and thalamic areas, we confirmed that single-neuron timescales correlate with anatomical hierarchy in the cortex but not in the thalamus^7^. In humans, timescales measured from the blood-oxygen-level-dependent (BOLD) signal follow established functional gradients in the striatum, thalamus, cerebellum, and hippocampus^15^. Within the thalamus, BOLD timescales specifically follow a ventrolateral-to-dorsomedial gradient, corresponding to the approximate locations of first-order and higherorder nuclei^15^. Thus, single-neuron timescales may likewise follow the anatomical layout of brain areas beyond the cortex, although discrepancies may arise due to differences in species (human versus mouse) and data modality (single-cell spiking versus BOLD)^3^. In addition, since BOLD reflects spatially averaged activity, denser sampling of individual neurons may be required to uncover the spatial and functional gradients of single-neuron timescales in the thalamus and other subcortical structures.

We found that spatial patterns of gene expression and genetically defined cell types predict variations in intrinsic timescales at a resolution finer than brain-area boundaries. Our results extend prior findings that gene expression profiles predict differences in timescales across cortical areas^9, 11^, by broadening the scope to the whole brain and sharpening the spatial resolution. Nonetheless, the predictive accuracy of our analysis was limited by reliance on anatomical features coarse-grained in 200 *µ*m voxels rather than cell-by-cell correspondences. Combining single-cell activity recordings with single-cell RNA sequencing^65^ may uncover even stronger relationships between gene expression and timescales. As genes affecting synaptic and membrane time constants are the strongest predictors of timescales^11^, cellular mechanisms such as slow synaptic currents^66–68^ or plateau potentials mediated by neuromodulators^69^ are likely among major contributors to sustaining persistent firing and generating long timescales.

The brain-wide span of our dataset revealed a universal power-law distribution of slow timescales across neurons, with an exponent near 2. This universality was evident in consistent scaling exponents obtained from neurons grouped by anatomical region, brain structure, recording session, or pooled across the entire brain. In primates, timescales in parietal, cingulate, and prefrontal cortices similarly follow a power-law distribution with an exponent near 2^51^, suggesting that the universal scaling is conserved across species and may provide a distributed, flexible neural substrate for cognitive computations over a broad range of timescales^51^. In contrast, fast timescales—below the power-law cutoff—dominated the regional variation in median timescales, as they contributed roughly half of the timescale distribution within each region. These fast timescales may therefore reflect the functional specialization of brain regions defined by their unique anatomical features. This multiscale architecture reconciles regional variations in timescales with the universality of a shared dynamical regime across the brain.

Our network models suggest that the universal scaling can emerge from common connectivity statistics across brain areas. Linear networks with fixed-point dynamics predict that the power-law exponent is determined by the shape of the eigenvalue distribution near zero for non-heavy-tailed connectivity. In contrast, nonlinear networks with chaotic dynamics predict that the exponent is set by the scaling of heavy-tailed connectivity. While linear networks require fine-tuning of synaptic gain to the edge of instability, nonlinear networks robustly generate the same power-law distribution across a broad range of synaptic gain values. The separation of fast and slow timescales, with slow timescales attributed to dynamics at the edge of instability or chaos, aligns with observations that cortical population activity exhibits a spectrum of timescales across distinct subspaces, some operating in subcritical and others in critical regimes^70, 71^. Cellular-resolution reconstructions of the mouse brain connectome^72, 73^ will reveal whether, and which, connectivity statistics are shared across brain areas, constraining network mechanisms for universal scaling.

Although our models are undoubtedly a stark oversimplification of brain circuitry, such abstractions may help distill general mechanisms invariant to implementation details. In particular, linear networks at the edge of instability also account for strong correlations between neurons^70^ and the power-law eigenspectrum of these correlations^74,75^, as observed in cortical data^76–81^. Nevertheless, our models lack biological realism, as they omit distinct cell types^23^, synaptic dynamics ^23,67,82^, and non-random structure in local^33,50,80,83–86^ and long-range inter-area connectivity^87,88^, all of which influence the timescales of single neurons. Incorporating these biological features into whole-brain models, enabled by emerging anatomical^8,43,44,89^ and activity datasets^31^ spanning the entire brain, will help establish a unified theoretical framework for linking structure, dynamics, and behavior at the scale of the whole brain.

## Methods

### Behavior and electrophysiology

We analyzed a dataset of brain-wide electrophysiological recordings of neural activity in mice collected by IBL^31^ (version 6^90^, https://doi.org/10.6084/m9.figshare.21400815.v6). This dataset consists of 354 Neuropixels recording sessions from 115 mice (C57BL/6; 80 male and 35 female, obtained from either Jackson Laboratory or Charles River). Mice were aged 13 − 122 weeks (mean 34.43 weeks, median 26.0 weeks) and weighed 16.1 − 36.2 g (mean 24.17 g, median 24.0 g) on the day of electrophysiological recording. The experiments and data collection were performed in 12 laboratories across Europe and the USA. A detailed description of the behavioral task and electrophysiology recordings can be found in the previous study^31^.

All experimental procedures involving animals were conducted in accordance with local laws and approved by the relevant institutional ethics committees. Approvals were granted by the Animal Welfare Ethical Review Body of University College London, under licences P1DB285D8, PCC4A4ECE, and PD867676F, issued by the UK Home Office. Experiments conducted at Princeton University were approved under licence 1876-20 by the Institutional Animal Care and Use Committee (IACUC). At Cold Spring Harbor Laboratory, approvals were granted under licences 1411117 and 19.5 by the institutional IACUC. The University of California at Los Angeles granted approval through IACUC licence 2020-121-TR-00. Additional approvals were obtained from the University Animal Welfare Committee of New York University (licence 18-1502); the IACUC at the University of Washington (licence 4461-01); the IACUC at the University of California, Berkeley (licence AUP-2016-06-8860-1); and the Portuguese Veterinary General Board (DGAV) for experiments conducted at the Champalimaud Foundation (licence 0421/0000/0000/2019).

Each session included at least 400 consecutive trials of the IBL decision-making task^31^, followed by a 10-minute passive period, which we used to estimate intrinsic timescales. During this passive period, head-fixed mice remained in the rig in a dark environment, engaging in spontaneous movements without external stimuli or rewards. We treated neural activity during the passive period as a proxy for spontaneous activity to estimate intrinsic neural timescales.

To examine how intrinsic timescales relate to task-variable encoding (stimulus, choice, and reward), we also analyzed neural activity during the task to assess selectivity of each neuron for these variables. On each trial of the task, a visual stimulus (Gabor patch) was presented on the left or right side (±35^*°*^ azimuth) of the screen. The contrast of the stimulus was randomly selected from a predefined set (100%, 25%, 12.5%, 6%, or 0%), and varied across trials. The probability of stimulus appearance on the left/right side was constant over a block of trials, at 20/80% (right block) or 80/20% (left block). Each block lasted for between 20 and 100 trials. To initiate a response, the mouse had to rotate a wheel to shift the stimulus to the center of the screen within a 60 s window. A response was recorded when the stimulus center crossed the 35^*°*^ azimuth line from its initial position. If the mouse correctly shifted the stimulus 35^*°*^ toward the center of the screen, it was immediately rewarded with 3 *µ*L of water. Conversely, if the mouse moved the stimulus 35^*°*^ away from the center, a timeout was triggered. In cases of incorrect responses or failure to reach either threshold within the 60 s window, a 500 ms burst of white noise was played, and the inter-trial interval was set to 2 s. The next trial began after a delay, followed by a quiescence period of 400–700 ms during which the mice had to hold the wheel still.

The electrophysiological data were recorded by at most two Neuropixels probes simultaneously during each session. The insertion locations of probes followed a grid covering the left hemisphere of the forebrain and midbrain and the right hemisphere of the cerebellum and hindbrain. In total, 547 insertions were performed, which yielded 295,501 neurons by performing spike-sorting (Kilosort) on the recordings, covering 267 brain areas. Next, quality control metrics were applied and led to 32,766 single units. Among 547 insertions, 504 insertions contained electrophysiological recordings during both task and passive periods of a session. Thus, we analyzed data from these 504 insertions exclusively, which included 27,960 single units spanning 223 brain regions.

### Estimation of intrinsic timescales

We estimated intrinsic timescales of single-neuron activity from the spike-count autocorrelation computed during 10-minute recordings of spontaneous activity at the end of each session. We computed the autocorrelation function AC(*t*_*j*_) at time lag *t*_*j*_ using spike counts 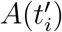 in Δ*t* = 5 ms bins^4^

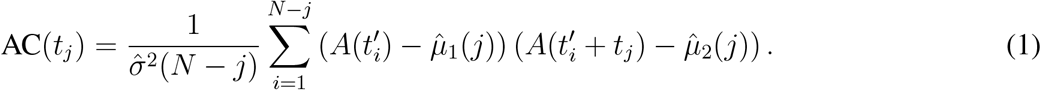

Here 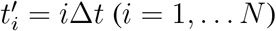 indicates time. The sample variance 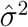 is

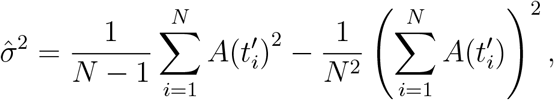

and 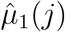 and 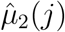 are two samples means for the original and shifted time series:

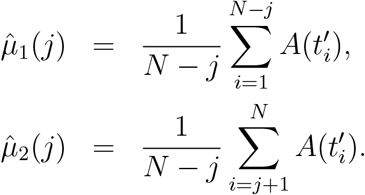

We directly fitted the autocorrelation with exponential decay functions, since the bias in autocorrelation estimates is negligible given the long duration of our time-series data^4^. We fitted the autocorrelation shape up to 5 s time lag to avoid overfitting noise in the autocorrelation tail^4^. We began fitting at the first time lag, since the drop in autocorrelation between zero and the first time lag reflects spiking irregularity rather than underlying dynamics^4, 91^. Some neurons also showed refractory-like behavior at the initial time lags, which we excluded from model fitting. Specifically, rather than fitting from the first time lag, we identified the earliest lag at which the autocorrelation began to decay and fitted the model using lags from that point onward^12, 19^.

To estimate the number and values of timescales for each neuron, we fitted the autocorrelation shape with linear mixtures of exponential decay functions:

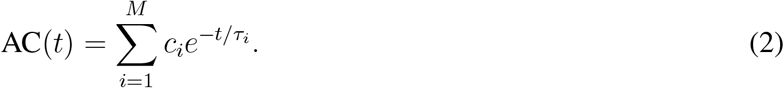

Here *τ*_*i*_ are the timescales, *c*_*i*_ are their mixing coefficients, and *M* is the number of distinct timescales. We fitted models with *M* ∈ 1, … 4 ranging from 1 (single timescale) up to 4 (four timescales) and selected the best-fitting model using the Bayesian information criterion (BIC). In addition, we required each timescale *τ*_*i*_ to contribute at least 1% to the overall autocorrelation shape 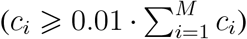 otherwise, we selected the model with fewer timescales, even if a more complex model was preferred by the BIC. This additional constraint ensured that our analysis focused only on timescales that substantially contributed to neural dynamics.

The model provided a good fit (coefficient of determination *R*^2^ *>* 0.5) for 11,468 out of 27,960 single units (41%), which were included in subsequent timescale analyses. This proportion of well-fitted neurons is consistent with previous studies of intrinsic timescales in single neurons^19, 22^. For the remaining neurons, the autocorrelation shape was either dominated by noise or oscillatory features that cannot be captured by exponential decay functions (Supplementary Fig. 1).

For each neuron, we defined the effective timescale as the average of individual timescales *τ*_*i*_ weighted by their mixing coefficients *c*_*i*_:

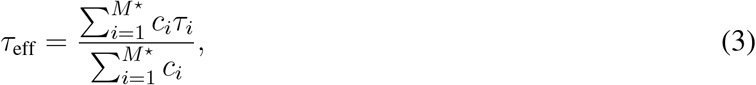

where *M*^*⋆*^ is the number of timescales in the selected model. The effective timescale provides a single summary statistic of the temporal integration properties of each neuron, enabling direct comparison across neurons with different numbers of fitted timescales. The effective timescale has a clear interpretation: the area under the original multi-timescale autocorrelation curve is equal to the area under a single-exponential decay with timescale *τ*_eff_. Thus, *τ*_eff_ captures the overall temporal persistence of neural activity, integrating over the multiple timescales that contribute to a neuron’s dynamics.

To facilitate comparison with previous studies of neural timescales, we also fitted the activity of each neuron with a single exponential function. All analyses based on this single-exponential fit included all neurons, without applying any selection criteria related to fit quality.

### Selectivity to task variables

To test the relationship between intrinsic timescales and encoding of task variables, we quantified the selectivity of single neurons to task variables: visual stimulus (left versus right location of the visual stimulus), choice (left versus right direction of wheel turning), and feedback (reward versus no reward) during the IBL decision-making task.

To quantify selectivity of single neurons to these three task variables, we used a nonparametric statistical test, the condition-combined Mann-Whitney U statistic, as previously described^31^. This method assigns a p-value for the selectivity of each neuron to a single task variable by comparing the sum of spike counts within a fixed time window between trials with two different values of that task variable, termed *V*_1_ (e.g., trials with stimulus on the left side) and *V*_2_ (e.g., trials with stimulus on the right side), respectively. For each variable, we used spike counts in time windows aligned to the relevant task event: [0, 100] ms from stimulus onset for the visual stimulus, [−100, 0] ms from first movement onset for choice, and [0, 200] ms from feedback onset for the reward.

To calculate the U statistic, we first computed spike counts for each trial, sorted them in ascending order, and assigned numeric ranks to the resulting array. We then computed the sum of ranks *R*_1_ and *R*_2_ for the spike counts in *n*_1_ and *n*_2_ trials associated with the task-variable values *V*_1_ and *V*_2_, respectively. The U statistic is given by

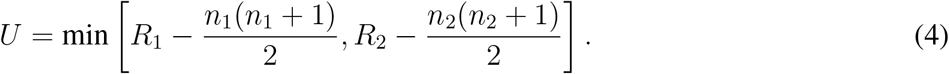

The probability of observing a difference in spike counts on *V*_1_ trials and those on *V*_2_ trials is defined as

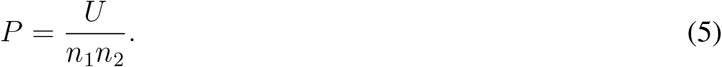

Since neural activity can be modulated by several task variables, we further designed a procedure to isolate the influence of individual task variables on neural activity. Specifically, for one task variable, we compared the spike counts from trials with this task-variable value *V*_1_ to those with the other value *V*_2_ and computed the U statistic, while holding the values of the other two task variables fixed. For example, we compared trials with the left versus the right side of the visual stimulus, while fixing choice and reward values. Following the same procedure, we analyzed other conditions where choice and reward variables have different fixed values.

The overall probability of identifying a spike-count difference in trials with different values of one task variable is given by a combination of the U statistic, *U*_*j*_, in each trial condition *j*^35^:

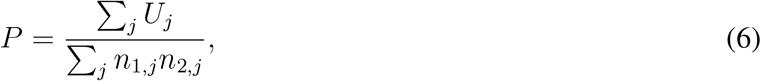

where *n*_1,*j*_ and *n*_2,*j*_ are the numbers of *V*_1_ and *V*_2_ trials, respectively, within the trial condition *j*.

To estimate the p-value of selectivity for one task variable, we used a permutation test. We randomly permuted trial labels for one task variable 3,000 times within each subset of trials with fixed values of all other task variables and then computed Mann-Whitney U statistic for each permutation. Since the block identity (left or right) shifts over time during each session, spurious correlations between neural activity and task variables can arise^45^. To avoid that, instead of a random permutation over the entire session, we randomly permuted trial labels with fixed values of all other task variables within each block, which effectively reduced the serial dependencies of task variables at the timescale of block duration. The p-value for each task variable is defined as the fraction of permutations where the statistic *P* is greater than in the data. We defined neurons with a p-value of less than 0.05 as being selective for this task variable and non-selective otherwise.

### Predicting timescales from anatomical features at fine spatial resolution

We tested whether anatomical features predict spatial variation in timescales at a resolution finer than brain-area boundaries. To this end, we modeled single-neuron timescales using ridge regression with predictors derived from either of two anatomical datasets: the Allen Gene Expression Atlas^44^ or the Allen Brain Cell Atlas^43^.

The Allen gene-expression atlas^44^ contains gene expression densities of 4,353 genes at each spatial location across the mouse brain, with a spatial resolution of 200 *µ*m. Since spatial expression densities are correlated across genes, we reduced the dimensionality of the gene expression data to minimize redundancy among predictors. We retained the top 275 principal components of the gene expression densities, which together captured 95% of the variance, and used them as predictors in the linear regression model.

The Allen Brain Cell Atlas^43^ provides cell-type annotations for 8.9 million single neurons across the whole adult mouse brain. Cell types were defined by clustering gene-expression profiles of all cells into 5,322 clusters, which were then grouped into 338 subclasses using 534 transcription factor marker genes. These 338 subclasses formed the set of cell types that we analyzed. From this dataset, we derived spatial densities of these 338 genetically defined cell types by computing the average number of cells of each type within voxels measuring 200 *µ*m per side. We then used these spatial densities as predictors in the linear regression model.

To fit the linear regression, we registered the timescale data to the same spatial resolution as the anatomical data. We assigned each single neuron to a voxel *i* and denoted its timescale as *τ*_*im*_, where 1 ⩽ *m* ⩽ *n*_*i*_ indexes the neurons and *n*_*i*_ is the number of neurons within voxel *i*. For each timescale *τ*_*im*_, the regression model predicted its value as a linear combination of voxel-specific predictors 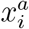, where *a* indexes predictors: 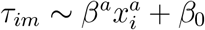. The coefficients *β*^*a*^ are the fitted parameters of the regression model, with *β*_0_ denoting the intercept.

As a baseline, we compared the performance of regression models based on anatomical features with a model that used only the brain region where each neuron was located. In this baseline model, the predictors 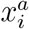 were binary indicator variables for a neuron’s brain region, with *a* = 1, 2, 3, …, 223 indexing regions defined in the Allen Common Coordinate Framework^31, 46^. For each brain region *a*, if voxel *i* belongs to this region, then 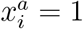, otherwise 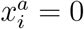.

We verified that the regression model predictions were not driven by spurious correlations between timescales and anatomical features, which could arise from independent spatial variation in both timescales and predictor variables across the brain^45^. To this end, we compared the performance of the regression models with that of surrogate models constructed using the phase randomization method to generate a null distribution of predictions^45^. Specifically, we applied a 3-dimensional Fourier transform to the spatial profiles of anatomical features, multiplied each Fourier coefficient by a random phase *e*^2*πiη*^, where *η* ∈ [0, 1] is sampled from a uniform distribution, and then applied the inverse Fourier transform to generate the surrogate spatial profiles. This procedure preserves the spatial power spectrum of the original data, thereby maintaining spatial correlation structure, while disrupting any relationship with the timescale data. We then computed *R*^2^ for regression models fitted with the surrogate predictors, repeating the procedure across 500 surrogate realizations, and compared these values with the *R*^2^ of the original model. For both gene-expression and cell-type density predictors, the original regression models predicted timescales significantly better than the surrogate models (*p <* 0.02, Supplementary Fig. 7).

To quantify the contribution of individual cell types to predicting timescales, we computed their unique explained variance. For each cell type, we generated a reduced model in which the spatial location (voxel index) of its density was randomly shuffled. We computed the difference in explained variance between the original and reduced models, Δ*R*^2^, as a measure of unique explained variance (Supplementary Fig. 8). Cell types with larger Δ*R*^2^ values had a greater impact on predicting timescales.

### Fitting power-law distribution of timescales

To evaluate whether the distribution of timescales was consistent with a power law and to estimate its exponent, we fitted a power-law model to the data using a standard maximum likelihood estimation method^48^. Since power-law scaling is generally expected only for sufficiently large timescales in the tail of the distribution, this method first determines an optimal cutoff value, restricting the power-law fit only to timescales above the cutoff. Starting from the smallest timescale, the algorithm increases the cutoff to each successive data point in ascending order and, for each value, computes the Kolmogorov–Smirnov (KS) distance between the data above the cutoff and the fitted power law. The cutoff that minimizes this KS distance was then selected and used in all subsequent analyses. No upper cutoff was imposed, so all data above the chosen lower cutoff were included in the fit.

To assess whether the timescale distribution was consistent with a power law, we performed a bootstrap-based statistical test that estimates the p-value for rejecting the power-law hypothesis^47^. We fitted the timescale distribution with a power-law model, estimating the optimal exponent *γ*, the lower cutoff *θ*, and the number of data points above the cutoff *N*_*θ*_, and computed the KS distance between the empirical data above the cutoff and the fitted truncated power-law distribution. We then generated an ensemble of *n* = 10, 000 synthetic datasets, each with *N*_*θ*_ samples drawn from a power-law distribution with exponent *γ*, discarding and resampling any points below the cutoff *θ* until all *N*_*θ*_ points satisfied *x* ⩾ *θ*. Each synthetic dataset was then independently fitted to a power-law model, producing an estimated exponent and the KS distance relative to its corresponding best-fitting distribution. This procedure generates the distribution of KS distances expected for a genuine power-law data. The p-value for rejecting the power-law hypothesis was defined as the fraction of synthetic datasets for which the KS distance exceeded that of the real data. A p-value smaller than 0.05 would indicate rejection of the power-law hypothesis. Following previous work, we adopted a conservative threshold of 0.1, such that p-values above this value imply that the power-law hypothesis cannot be ruled out^47^.

In all major brain structures, as well as in the data pooled from all regions, the p-values exceeded the 0.1 threshold (Supplementary Table 3), indicating that the timescales were consistent with a power-law distribution. Since different brain structures exhibited distinct power-law cutoff values (Fig. 5d), we used the maximum observed cutoff (Pons, Supplementary Table 3) when pooling data across all brain structures, ensuring that the remaining data were consistent with a power-law distribution in each structure.

We additionally compared the goodness of fit of the power-law model against the exponential and log-normal alternatives. For each dataset, we fitted all three models to the data above the power-law cutoff using standard maximum likelihood estimation^48^. The log-likelihood ratio (LLR) for pairwise comparison between the alternative models is

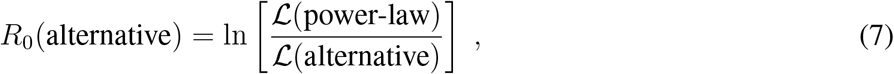

where L(power-law) and L(alternative) denote the likelihoods of the power-law and the alternative model fits, respectively. For computing significance, we used a normalized LLR^47^:

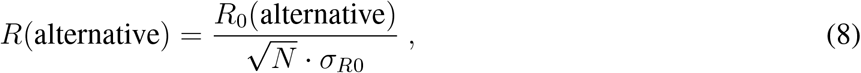

where *N* is the number of data points included in the analysis, *σR*_0_ is the standard deviation of the loglikelihood ratio across all data points. All fitting procedures and model comparisons were performed using the Python package powerlaw, designed for the analysis of heavy-tailed distributions^48^.

In all major brain structures, as well as in data pooled across all regions, the LLR comparing power-law and exponential models was significantly positive (Supplementary Table 5), confirming that timescales followed a heavy-tailed distribution. In all brain structures and in the pooled dataset, the LLR comparing power-law and log-normal models was not significantly different from zero (Supplementary Table 5). To interpret this result, we simulated data sampled from a true power-law distribution with exponent *γ* = 2 and cutoff matching the empirical data, using *N*_*θ*_ = 1, 000 data points, larger than most empirical datasets we analyzed. In most cases, the resulting LLRs were negative (slightly favoring the log-normal) but not statistically significant, and only 2 out of 1,000 simulated datasets yielded a significant LLR, both favoring the power-law hypothesis. These simulations indicate that our observations are consistent with expectations for a true power-law distribution given the limited sample sizes.

### Distribution of timescales in linear networks

We derived analytical expressions for the distribution of timescales in a linear network model and examined how this distribution depends on the connectivity eigenvalue distribution, synaptic time constant, and synaptic gain parameter.

The dynamics of a linear network model with *N* units are governed by the equations:

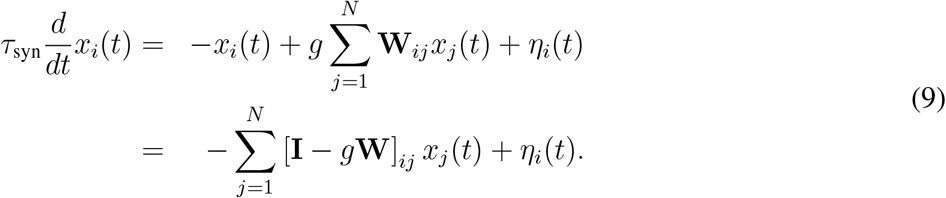

Here *x*_*i*_(*t*) denotes the activity of unit *i* (*i* = 1, … *N*), and *η*_*i*_(*t*) is a Gaussian white noise with ⟨*η*_*i*_(*t*)⟩ = 0 and ⟨*η*_*i*_(*t*)*η*_*j*_(*t*^*′*^)⟩ = *δ*(*t* − *t*^*′*^)*δ*_*ij*_*D*_*i*_. We assume that the noise magnitude is constant across units: *D*_*i*_ = *D*. The connectivity matrix is **W**_*ij*_. We define the effective connectivity matrix as **M**_*ij*_ = [**I** − **W**]_*ij*_. The parameter *τ*_syn_ denotes the synaptic time constant, and *g* is the synaptic gain parameter. To simplify notation, we first derive the timescale distributions in linear networks with *τ*_syn_ = 1 and *g* = 1, and subsequently examine the effects of these parameters on the timescale distribution.

First, we analyze how the connectivity matrix determines the distribution of timescales, setting *τ*_syn_ = 1 and *g* = 1. When the network activity fluctuates around a fixed point, the time-delayed correlation matrix between the activity of unit pairs *i* and *j* is defined by

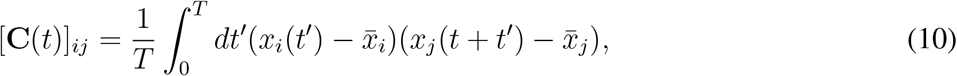

where *T* is the time window, and 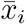 is the time-averaged activity of unit *i*.

To compute the timescales of activity autocorrelation, we diagonalize the effective connectivity matrix. The right-eigenvectors of matrix **M** are given by:

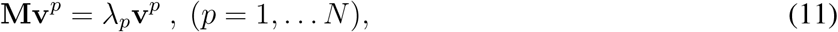

where *p* indexes the eigenvectors. Similarly, the left-eigenvectors of **M** are defined by

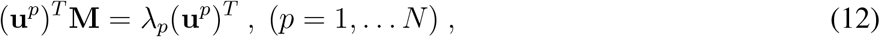

We then analytically derive the expression for the correlation matrix **C**(*t*)^92^:

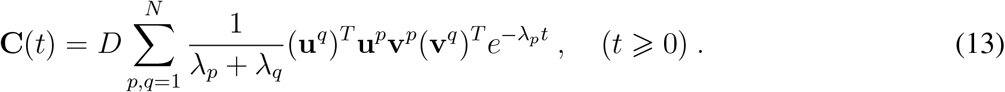

The autocorrelation of the activity of unit *i* is given by the diagonal element **C**_*ii*_. This autocorrelation is a linear combination of *N* eigenmodes, with timescales 1*/λ*_*p*_ (*p* = 1, … *N*) determined by the inverse eigenvalues of **M** (Eq. 13). In simulations, we found that the distribution of single-neuron timescales *τ*, estimated by fitting their autocorrelation with an exponential function, closely matches the distribution of the network timescales given by the inverse eigenvalues 1*/λ*_*p*_ (Supplementary Fig. 12). Therefore, we used the inverse eigenvalues 1*/λ*_*p*_ to simulate timescale distributions in linear networks (Fig. 6c,e,g).

The distribution of eigenvalues *λ* is denoted by *f* (*λ*), with the range *λ* ∈ [*λ*_min_, *λ*_max_]. For a stable fixed point, the smallest eigenvalue must be positive *λ*_min_ *>* 0, so the corresponding range of timescales is *τ* ∈ [1*/λ*_max_, 1*/λ*_min_]. To generate very slow timescales, *λ*_min_ must approach zero, *λ*_min_ ≈ 0. In the limit *λ*_min_ → 0, when the dynamics operate at the edge of instability, the maximal timescale diverges *τ* → ∞.

To obtain the distribution of timescales, we used the change-of-variable rule for a random variable and its reciprocal. Specifically, for a positive random variable *x* with probability density function *f* (*x*), the probability density function of its reciprocal *y* = 1*/x* is given by *g*(*y*) = (1*/y*^2^) · *f* (1*/y*). Accordingly, given an eigenvalue distribution *f* (*λ*), the timescale distribution *g*(*τ*) is given by

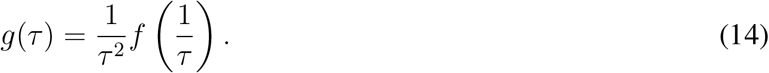

For long timescales, the distribution *g*(*τ*) is determined by the behavior of *f* (*λ*) at small *λ*. We consider the case where *f* (*λ*) can be approximated by a power function *f* (*λ*) ≈ *λ*^*ζ*^, for small *λ* below a cutoff value *λ*_cutoff_. In this regime, timescales exceeding *τ*_cutoff_ = 1*/λ*_cutoff_ follow a power-law distribution with exponent

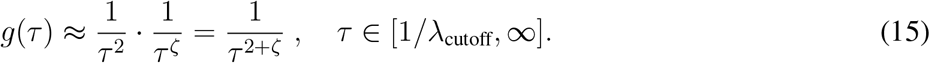

Thus, we found that in linear networks set at the edge of instability, the timescale distribution can follow a power law with an exponent determined by the shape of the connectivity eigenvalue distribution near zero.

This theory enabled us to systematically construct networks that generate a power-law distribution of timescales with tunable exponents by appropriately shaping the connectivity eigenvalue distribution. We illustrate this theory using example linear networks with different connectivity eigenvalue distributions and the corresponding timescale exponents.

### Uniform distribution

For a uniform eigenvalue distribution, *f* (*λ*) = const, hence *ζ* = 0 for the entire range of eigenvalues. Therefore, the entire range of timescales follows a power law with exponent *γ* = 2^51^.

### Semicircle distribution

For a semicircle distribution of eigenvalues

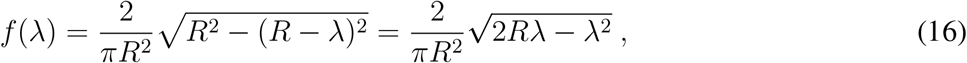

where *R* is the radius parameter. For small *λ* such that 2*Rλ* ≫ *λ*^2^, we retain only the leading term in the square root to obtain 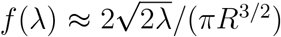. Hence, *ζ* = 0.5, and long timescales follow a power law with exponent *γ* = 2.5. We estimate the cutoff eigenvalue by setting 2*Rλ* = *λ*^2^, resulting in *λ*_cutoff_ = 2*R*. The corresponding cutoff timescale is *τ*_cutoff_ ≈ 1*/*(2*R*).

### Gamma distribution

When the eigenvalues follow a gamma distribution with shape parameter *α* and scale parameter 1, their probability density is given by

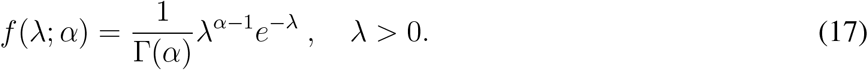

For small *λ*, the density is approximately *f* (*λ*; *α*) ≈ *λ*^*α−*1^*/*Γ(*α*). Hence, *ζ* = *α* − 1, and long timescales follow a power law with exponent *γ* = *α* + 1. We estimate the cutoff eigenvalue by setting *e*^*−λ*^ = 1*/*2, the point at which *e*^*−λ*^ substantially deviates from 1, resulting in *λ*_cutoff_ = ln(2). The corresponding cutoff timescale is *τ*_cutoff_ ≈ 1*/* ln(2).

Next, we analyze how the timescale distribution depends on synaptic time constant *τ*_syn_ and synaptic gain *g* parameters (Eq. 9). First, the synaptic time constant *τ*_syn_ directly affects the magnitude of timescales. Specifically, dividing both sides of Eq. 9 by *τ*_syn_ shows that the synaptic time constant rescales the effective connectivity by a factor of 1*/τ*_syn_. Consequently, all eigenvalues *λ* of the effective connectivity are uniformly scaled by 1*/τ*_syn_, and all timescales *τ* are proportionally scaled by *τ*_syn_. Thus, both the cutoff timescale and the median timescale are rescaled by *τ*_syn_. Second, increasing the synaptic gain *g* reduces the distance to the instability threshold. Specifically, the boundary of the eigenvalue distribution *f* (*λ*) is determined by the minimal eigenvalue of the effective connectivity matrix **I** − *g***W**. This minial eigenvalue is given by *λ*_min_ = 1 − *gλ*_*w*,max_, where *λ*_*w*,max_ *>* 0 is the maximal eigenvalue of **W**. Thus, *g* directly controls the magnitude of *λ*_min_, which quantifies the distance to the instability threshold at 0. As *λ*_min_ increases, the longest timescale becomes bounded, leading to a collapse of the power-law timescale distribution.

### Distribution of timescales in nonlinear networks

We simulated a nonlinear recurrent network model to investigate how timescales of single units depend on the connectivity distribution, synaptic time constant, and synaptic gain parameter.

The dynamics of a nonlinear network model with *N* units are governed by the equations:

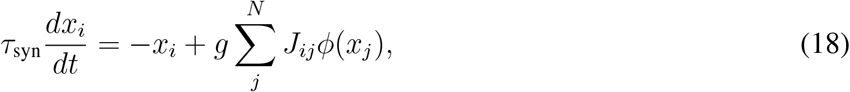

where *ϕ*(*x*) = tanh(*x*) is a single-unit activation function, *x*_*i*_ is the activity of unit *i, g* is the synaptic gain parameter, and *τ*_syn_ represents the synaptic time constant. Each element of the connectivity matrix *J*_*ij*_ is an independent random variable sampled from an *α*-stable distribution 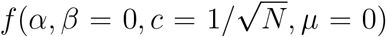, where 0 *< α <* 2. The *α*-stable distribution has an asymptotic power-law tail with exponent *γ*_*c*_ = 1 + *α*, such that Prob(*J*_*ij*_ = *x*) ∝ 1*/*|*x*|^1+*α*^ as *x* → ±∞. When the synaptic gain *g* exceeds a critical value *g*_crit_, the network activity displays chaotic dynamics, and the timescales of single-unit autocorrelations exhibit a heavy-tailed distribution^52^.

We simulated nonlinear networks with 1,000 units for varying values of parameters *α, τ*_syn_, and *g*. For each set of parameters, we performed 100 simulations with random initial conditions and random realizations of connectivity weights sampled from an *α*-stable distribution. Each simulation was run for 14,400 time steps. In each simulation, we estimated timescales of single units by fitting their activity autocorrelation with an exponential function. We then pooled timescales from all simulations into a single distribution.

## Data availability

The IBL dataset is publicly available on Figshare at https://doi.org/10.6084/m9.figshare. 21400815.v6. Instructions for downloading the data are available on the IBL GitHub at https://int-brain-lab.github.io/iblenv/notebooks_external/data_release_brainwidemap.html. The dataset can be explored and visualized athttp://viz.internationalbrainlab.org.

## Code availability

The source code to reproduce the results of this study will be made publicly available on GitHub upon publication.

## Acknowledgements

This work was supported by the National Institutes of Health grants 1U19NS123716 (YS, TAE), the Simons Foundation (YS, TAE), Wellcome Trust grants 209558 and 216324 (IBL), the German Research Foundation (DFG) through Germany’s Excellence Strategy (EXC-Number 2064/1, PN 390727645) (RZ), Max Planck Society (RZ), the German Federal Ministry of Education and Research (Tübingen AI Center, FKZ: 01IS18039) (AL), the Sofja Kovalevskaja Award from the Alexander von Humboldt Foundation endowed by the Federal Ministry of Education and Research (RZ, AL). We thank the International Max Planck Research School for the Mechanisms of Mental Function and Dysfunction (IMPRS-MMFD) and the Joachim Herz Foundation for their support. We thank Braden Brinkman, William Bialek, David Dahmen, James Fitzgerald, Pulin Gong, Brendan Harris, Tilo Schwalger, and Xiao-Jing Wang for discussions and valuable suggestions. We thank Matteo Carandini, Kenneth Harris, and Nicholas Steinmetz for constructive feedback on the project and manuscript.

## Author Contributions

YS, RZ, AL, and TAE designed the study. YS and RZ performed the research. YS, RZ, AL, and TAE discussed the findings and wrote the paper.

## Competing interests

The authors declare no competing interests.

## Notes

### Competing Interest Statement

The authors have declared no competing interest.

