## Supplementary Information for "Brain-wide organization of intrinsic timescales at single-neuron resolution"

### Supplementary Materials for: Brain-wide organization of intrinsic timescales at single-neuron resolution

### Contents

**1 Supplementary Tables** **2**

**2 Supplementary Figures** **7**

### 1 Supplementary Tables

| Cortical region ID | Acronym | Region name |
|---|---|---|
| 39 | ACAd | Anterior cingulate area dorsal part |
| 44 | ILA | Infralimbic area |
| 48 | ACAv | Anterior cingulate area ventral part |
| 104 | AId | Agranular insular area dorsal part |
| 111 | AIp | Agranular insular area posterior part |
| 119 | AIv | Agranular insular area ventral part |
| 184 | FRP | Frontal pole cerebral cortex |
| 329 | SSp-bfd | Primary somatosensory area barrel field |
| 337 | SSp-ll | Primary somatosensory area lower limb |
| 345 | SSp-m | Primary somatosensory area mouth |
| 353 | SSp-n | Primary somatosensory area nose |
| 361 | SSp-tr | Primary somatosensory area trunk |
| 369 | SSp-ul | Primary somatosensory area upper limb |
| 378 | SSs | Supplemental somatosensory area |
| 385 | VISp | Primary visual area |
| 394 | VISam | Anteromedial visual area |
| 409 | VISl | Lateral visual area |
| 417 | VISrl | Rostrolateral visual area |
| 425 | VISpl | Posterolateral visual area |
| 533 | VISpm | Posteromedial visual area |
| 541 | TEa | Temporal association areas |
| 723 | ORBl | Orbital area lateral part |
| 731 | ORBm | Orbital area medial part |
| 746 | ORBvl | Orbital area ventrolateral part |
| 879 | RSPd | Retrosplenial area dorsal part |
| 886 | RSPv | Retrosplenial area ventral part |
| 894 | RSPagl | Retrosplenial area lateral agranular part |
| 972 | PL | Prelimbic area |
| 985 | MOp | Primary motor area |
| 993 | MOs | Secondary motor area |
| 1002 | AUDp | Primary auditory area |
| 1011 | AUDd | Dorsal auditory area |
| 1027 | AUDpo | Posterior auditory area |
| 312782546 | VISa | Anterior area |
| 312782574 | VISli | Laterointermediate area |
| 312782628 | VISpor | Postrhinal area |

**Supplementary Table 1. Cortical regions included in the analysis of the relationship between timescales and anatomical hierarchy scores.** For each region, the table lists the brain region ID based on the Allen Common Coordinate Framework, its acronym, and full name [1].

| Thalamic region ID | Acronym | Region name |
|---|---|---|
| 127 | AM | Anteromedial nucleus |
| 149 | PVT | Paraventricular nucleus of the thalamus |
| 155 | LD | Lateral dorsal nucleus of thalamus |
| 170 | LGd | Dorsal part of the lateral geniculate complex |
| 181 | RE | Nucleus of reuniens |
| 218 | LP | Lateral posterior nucleus of the thalamus |
| 255 | AV | Anteroventral nucleus of thalamus |
| 362 | MD | Mediodorsal nucleus of thalamus |
| 366 | SMT | Submedial nucleus of the thalamus |
| 475 | MG | Medial geniculate complex |
| 575 | CL | Central lateral nucleus of the thalamus |
| 599 | CM | Central medial nucleus of the thalamus |
| 629 | VAL | Ventral anterior-lateral complex of the thalamus |
| 685 | VM | Ventral medial nucleus of the thalamus |
| 718 | VPL | Ventral posterolateral nucleus of the thalamus |
| 733 | VPM | Ventral posteromedial nucleus of the thalamus |
| 907 | PCN | Paracentral nucleus |
| 930 | PF | Parafascicular nucleus |
| 1020 | PO | Posterior complex of the thalamus |
| 1029 | POL | Posterior limiting nucleus of the thalamus |
| 1113 | IAD | Interanterodorsal nucleus of the thalamus |
| 560581563 | PIL | Posterior intralaminar thalamic nucleus |

**Supplementary Table 2. Thalamic regions included in the analysis of the relationship between timescales and anatomical hierarchy scores.** For each region, the table lists the brain region ID based on the Allen Common Coordinate Framework, its acronym, and full name [1].

| Brain structure | Acronym | $N$ | $N_\theta$ | Cutoff $\theta$ (s) | KS distance | p-value |
|---|---|---|---|---|---|---|
| All brain regions | | 10,279 | 1,641 | 1.2 | 0.023 | 0.12 |
| Isocortex | CTX | 2,439 | 553 | 0.7 | 0.026 | 0.29 |
| Olfactory areas | OLF | 468 | 158 | 0.2 | 0.035 | 0.71 |
| Hippocampal formation | HPF | 1,327 | 370 | 0.1 | 0.027 | 0.48 |
| Cortical subplate | CTXsp | 175 | 114 | 0.02 | 0.043 | 0.76 |
| Striatum | STR | 1,143 | 154 | 0.5 | 0.053 | 0.19 |
| Pallidum | PAL | 164 | 42 | 0.6 | 0.085 | 0.22 |
| Thalamus | TH | 2,601 | 463 | 0.4 | 0.024 | 0.50 |
| Hypothalamus | HY | 82 | 34 | 0.6 | 0.094 | 0.46 |
| Midbrain | MB | 914 | 286 | 1.1 | 0.027 | 0.66 |
| Pons | P | 215 | 93 | 1.2 | 0.048 | 0.76 |
| Medulla | MY | 446 | 246 | 0.6 | 0.045 | 0.14 |
| Cerebellum | CB | 305 | 143 | 0.8 | 0.046 | 0.39 |

**Supplementary Table 3. Bootstrap-based statistical test to assess whether the timescale distribution is consistent with a power law.** Results of the statistical test that estimates the p-value for rejecting the hypothesis that the timescales follow a power-law distribution in each major brain structure and in the data pooled from all regions. For each dataset, the table lists the number of neurons $N$, the number of neurons with timescales exceeding the optimal cutoff value $N_\theta$, the optimal cutoff value $\theta$, the KS distance to the best-fitting power law, and a p-value for rejecting the power-law hypothesis. A p-value greater than $0.1$ implies that the power-law hypothesis cannot be ruled out. In all brain structures and in the data pooled from all regions, the p-values exceeded the $0.1$ threshold, indicating that the timescales were consistent with a power-law distribution.

| Brain structure | Acronym | $N$ | $N_\theta$ | Cutoff $\theta$ (s) | KS distance | p-value |
|---|---|---|---|---|---|---|
| All brain regions | | 26,888 | 4,461 | 3.95 | 0.013 | 0.2201 |
| Isocortex | CTX | 6,489 | 770 | 5.47 | 0.024 | 0.5284 |
| Olfactory areas | OLF | 1,111 | 309 | 1.85 | 0.036 | 0.5702 |
| Hippocampal formation | HPF | 2,943 | 523 | 3.10 | 0.020 | 0.9377 |
| Cortical subplate | CTXsp | 498 | 104 | 2.86 | 0.073 | 0.3083 |
| Striatum | STR | 2,820 | 509 | 2.48 | 0.013 | 0.9998 |
| Pallidum | PAL | 357 | 64 | 3.57 | 0.070 | 0.6702 |
| Thalamus | TH | 4,794 | 771 | 3.67 | 0.019 | 0.8305 |
| Hypothalamus | HY | 270 | 105 | 2.24 | 0.049 | 0.8174 |
| Midbrain | MB | 2,043 | 568 | 3.13 | 0.025 | 0.6933 |
| Pons | P | 524 | 220 | 2.65 | 0.051 | 0.3198 |
| Medulla | MY | 787 | 253 | 2.37 | 0.044 | 0.4222 |
| Cerebellum | CB | 1,381 | 358 | 3.59 | 0.038 | 0.4001 |

**Supplementary Table 4. Bootstrap-based statistical test to assess whether timescales estimated by single-exponential fits are consistent with a power-law distribution.** Results of the statistical test that estimates the p-value for rejecting the hypothesis that the timescales follow a power-law distribution in each major brain structure and in the data pooled from all regions. For each dataset, the table lists the number of neurons $N$, the number of neurons with timescales exceeding the optimal cutoff value $N_\theta$, the optimal cutoff value $\theta$, the KS distance to the best-fitting power law, and a p-value for rejecting the power-law hypothesis. To compute the p-value, we generated an ensemble of $n = 100,000$ synthetic datasets. A p-value greater than $0.1$ implies that the power-law hypothesis cannot be ruled out. In all brain structures and in the data pooled from all regions, the p-values exceeded the $0.1$ threshold, indicating that the timescales were consistent with a power-law distribution. All neurons were included in this analysis.

| Brain structure | Acronym | $N$ | $N_\theta$ | $R$ (lognormal) | p(lognormal) | $R$ (exp) | p(exp) |
|---|---|---|---|---|---|---|---|
| All brain regions | | 10,279 | 1,641 | 0.031 | 0.54 | 1647.06 | $< 1.0 \times 10^{-10}$ |
| Isocortex | CTX | 2,439 | 553 | 0.002 | 0.98 | 649.85 | $< 1.0 \times 10^{-10}$ |
| Olfactory areas | OLF | 468 | 158 | 0.002 | 0.88 | 189.18 | $5.7 \times 10^{-10}$ |
| Hippocampal formation | HPF | 1,327 | 370 | $-0.68$ | 0.46 | 481.78 | $< 1.0 \times 10^{-10}$ |
| Cortical subplate | CTXsp | 175 | 114 | $-0.085$ | 0.81 | 171.79 | $7.1 \times 10^{-3}$ |
| Striatum | STR | 1,143 | 154 | 0.002 | 0.95 | 312.02 | $< 1.0 \times 10^{-10}$ |
| Pallidum | PAL | 164 | 42 | $-0.44$ | 0.55 | 10.62 | $1.5 \times 10^{-1}$ |
| Thalamus | TH | 2,601 | 463 | $-0.72$ | 0.37 | 710.41 | $< 1.0 \times 10^{-10}$ |
| Hypothalamus | HY | 82 | 34 | $-0.046$ | 0.37 | 25.44 | $9.0 \times 10^{-3}$ |
| Midbrain | MB | 914 | 286 | $-0.47$ | 0.48 | 209.82 | $< 1.0 \times 10^{-10}$ |
| Pons | P | 215 | 93 | $-0.20$ | 0.63 | 52.71 | $2.3 \times 10^{-7}$ |
| Medulla | MY | 446 | 246 | $-1.86$ | 0.14 | 164.08 | $< 1.0 \times 10^{-10}$ |
| Cerebellum | CB | 305 | 143 | $-1.36$ | 0.23 | 75.14 | $3.7 \times 10^{-8}$ |

**Supplementary Table 5. Comparison of power-law fits with exponential and log-normal alternatives.** Results of comparing the goodness of fit of the power-law model against the exponential and log-normal alternatives for the timescale distributions in each major brain structure and in the data pooled from all regions. For each dataset, the table lists the number of neurons $N$, the number of neurons with timescales exceeding the optimal cutoff value $N_\theta$, the normalized log-likelihood ratio (LLR) between the power-law and the lognormal model $R$(lognormal) and the corresponding p-value p(lognormal), and the normalized LLR between the power-law and exponential model $R$(exp) and the corresponding p-value p(exp). A positive LLR indicates that the power-law model provides a better fit than the alternative, whereas a negative LLR favors the alternative.

#### 2 Supplementary Figures

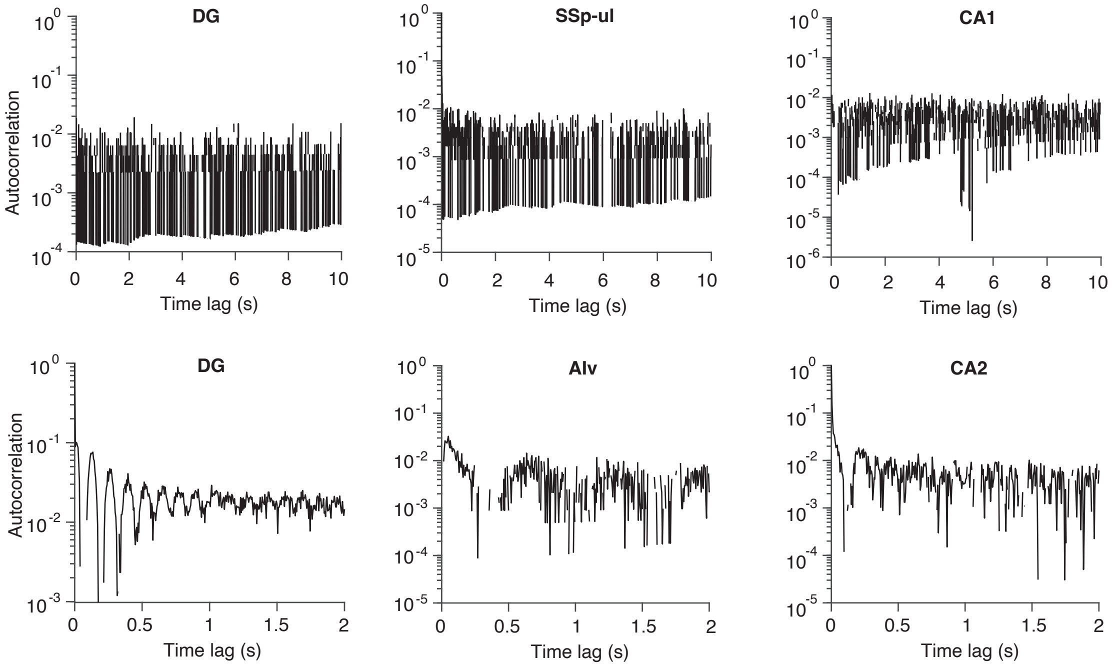


**Supplementary Figure 1. Example autocorrelations that cannot be adequately captured by exponential decay functions.** Autocorrelations of example single neurons poorly fitted by the exponential decay model (coefficient of determination $R^2 \leqslant 0.5$). The autocorrelation shapes were either dominated by noise (top row; DG: Dentate gyrus; SSpul: Primary somatosensory area upper limb; CA1: Field CA1) or oscillatory features (lower row; DG: Dentate gyrus; Alv: alveus; CA2: Field CA2).

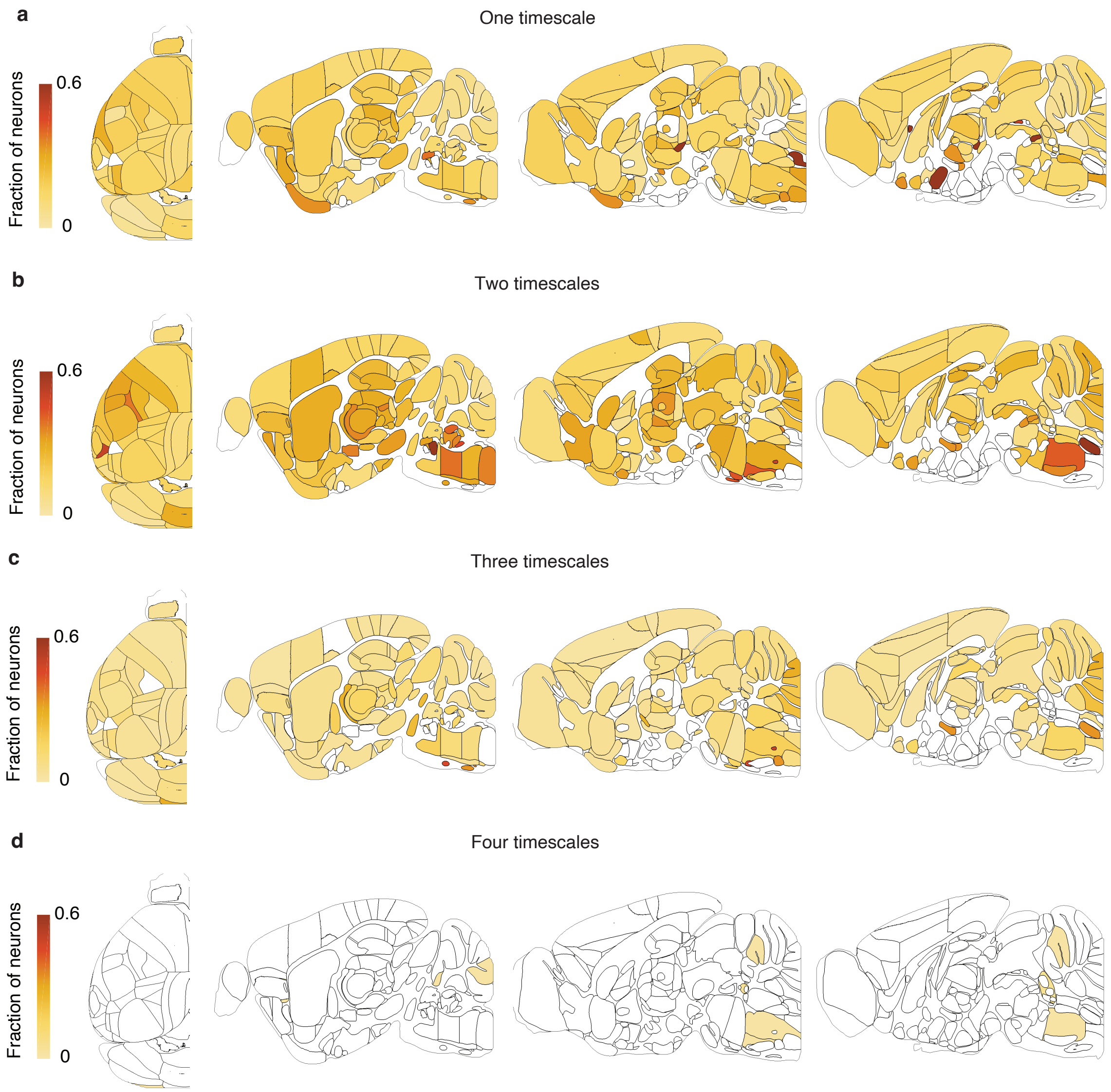


**Supplementary Figure 2. Distribution of neurons with different number of timescales across the brain.** **a**, Brain-slice plots (top and sagittal views) showing the fraction of neurons best fit by a single-timescale model in each brain region. **b**, Same as panel a for neurons best fit by a two-timescale model. **b**, Same as panel a for neurons best fit by a three-timescale model. **b**, Same as panel a for neurons best fit by a four-timescale model.

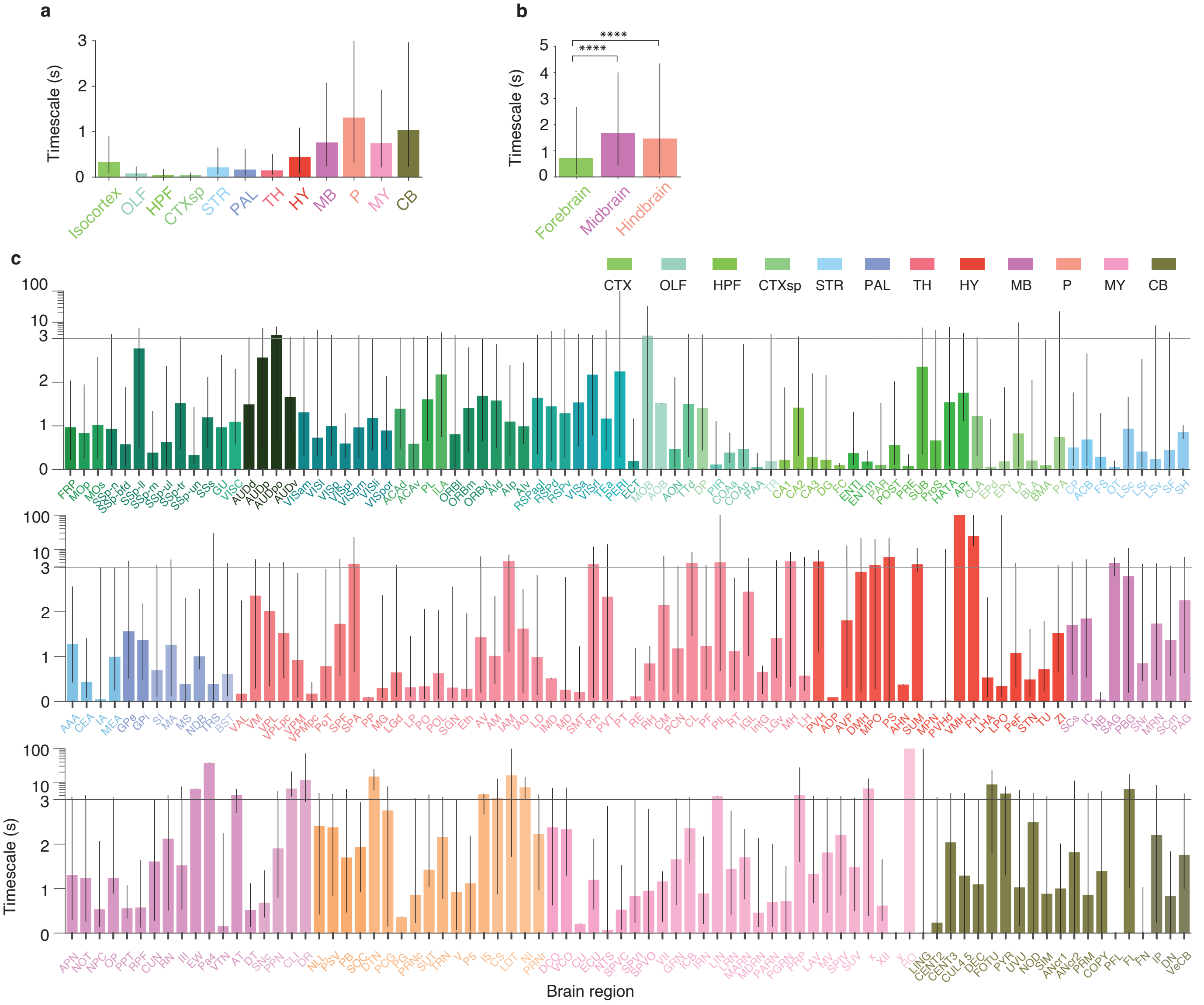


**Supplementary Figure 3. A brain-wide map of intrinsic timescales estimated from single-exponential fit. a**, Timescales in 12 major brain structures. Bar height indicates the median, and error bars represent the 25th to 75th percentiles. **b**, Timescales for the three major brain divisions: forebrain (including isocortex, olfactory areas, hippocampal formation, cortical subplate, striatum, pallidum, thalamus, and hypothalamus), midbrain, and hindbrain (including medulla, pons, and cerebellum). Bar height indicates the median (forebrain 0.72 s, $n = 19,979$ neurons; midbrain 01.66 s, $n = 2,144$; hindbrain 1.47 s, $n = 2,831$), and error bars represent the 25th to 75th percentiles. Timescales were significantly longer in the midbrain and hindbrain than in the forebrain (two-sided Wilcoxon rank sum test, $^{****}p < 10^{-5}$). **c**, Timescales for all brain regions across the 12 major brain structures indicated by color (CTX, isocortex; OLF, olfactory areas; HPF, hippocampal formation; CTXsp, cortical subplate; STR, striatum; PAL, pallidum; TH, thalamus; HY, hypothalamus; MB, midbrain; P, pons; MY, medulla; CB, cerebellum). Bar height indicates the median, and error bars represent the 25th to 75th percentiles of the timescale distribution. The y-axis transitions from a linear to logarithmic scale (horizontal line) to accommodate the wide distribution of timescales. All neurons were included in these analyses.

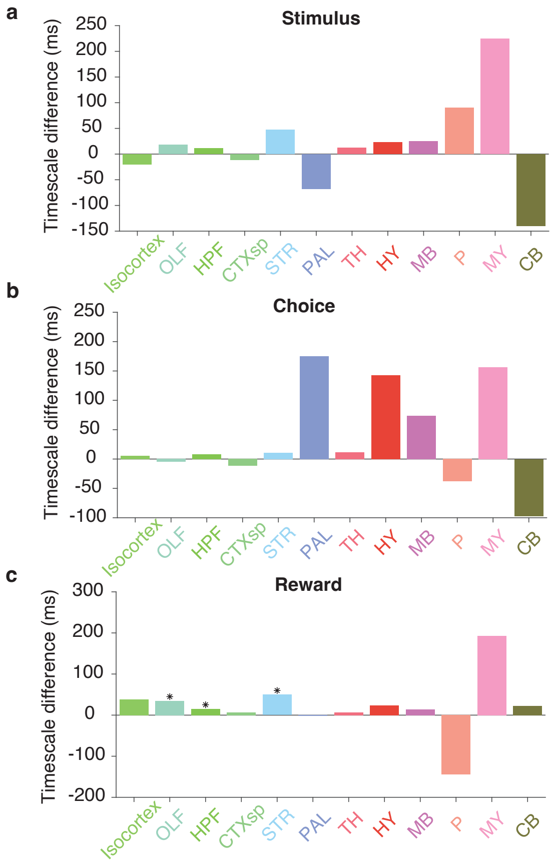


**Supplementary Figure 4. Differences in timescales between selective and non-selective neurons for each task variable.** **a**, Difference in median effective timescales between selective and non-selective neurons for the visual stimulus in each major brain structure. **b**, Same as panel a for choice. **c**, Same as panel a for reward.

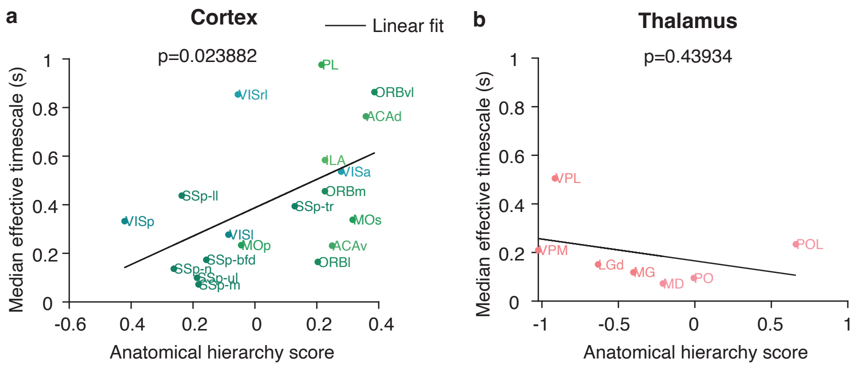


**Supplementary Figure 5. Relationship between anatomical hierarchy scores and effective timescales in previously studied regions.** **a**, Median effective timescales increased with the anatomical hierarchy scores (Pearson correlation coefficient, $r = 0.52$, $p = 0.02$, $n = 19$) in the subset of cortical areas studied previously [2]. Black line shows the linear fit. **b**, Median effective timescales did not significantly correlate with the anatomical hierarchy scores (Pearson correlation coefficient, $r = -0.35$, $p = 0.44$, $n = 7$) in the subset of thalamic areas studied previously [2]. Black line shows the linear fit.

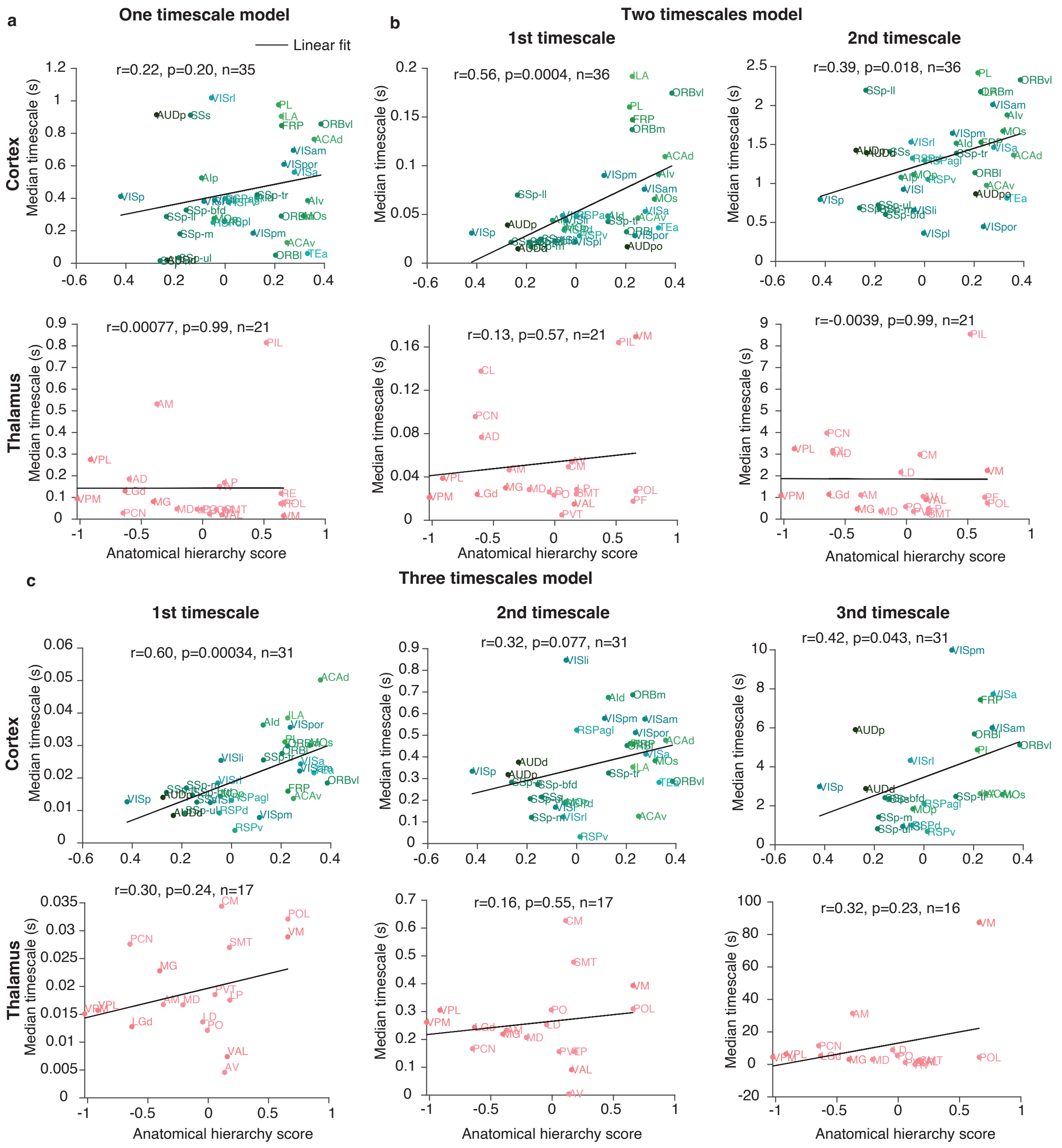


**Supplementary Figure 6. Relationship between anatomical hierarchy scores and individual fitted timescales of single neurons.** **a**, Relationship between anatomical hierarchy scores and median timescales for single-timescale neurons in cortex (top) and thalamus (lower panel). **b**, Relationship between anatomical hierarchy scores and median constituent timescales (1st and 2nd) for two-timescale neurons in cortex (top) and thalamus (lower panel). **b**, Same as panel b for three-timescale neurons. In all panels, $r$ denotes the Pearson correlation coefficient, $p$ the p-value, and $n$ the sample size. Black lines show the linear fit.

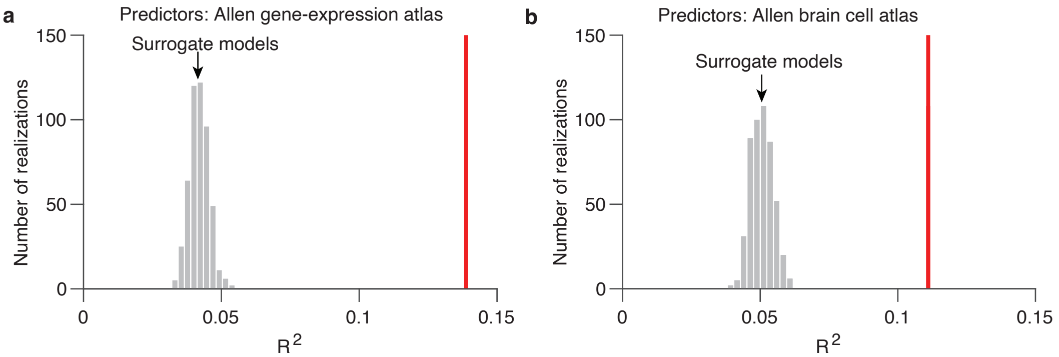


**Supplementary Figure 7. Phase-randomization test of models predicting timescales from anatomical features. a**, The prediction accuracy $R^2$ of regression models with predictors from the Allen gene-expression atlas (red line) was significantly higher than that of surrogate models based on phase randomization, which preserves the spatial correlation structure of predictors but eliminates their relationship to the timescale data (gray histogram, 500 simulations). **b**, Same as panel a for predictors from the Allen brain cell atlas.

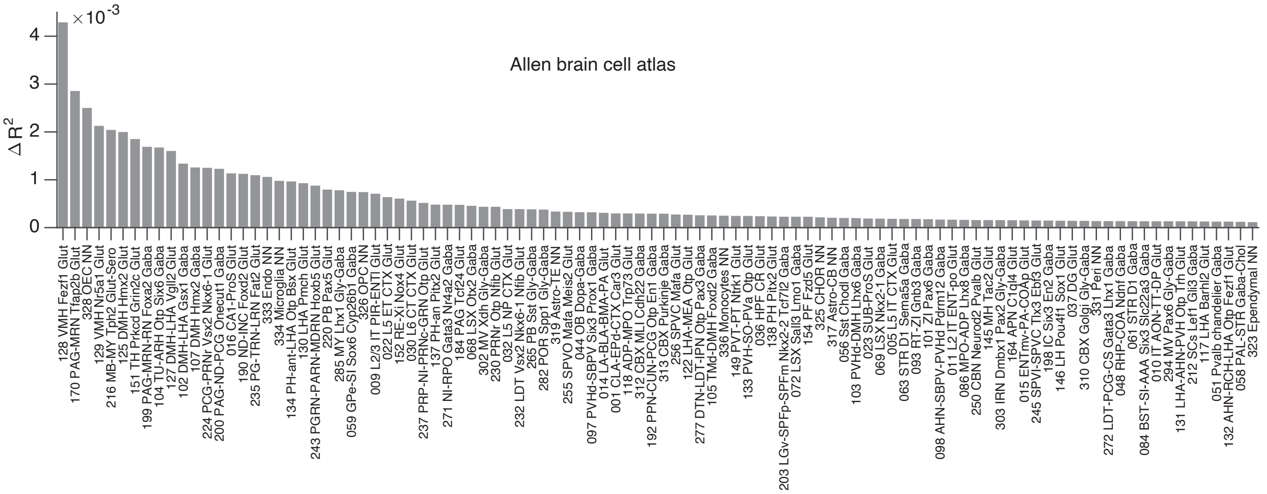


**Supplementary Figure 8. Contribution of individual cell types to predicting timescales.** Top 100 genetically defined cell types ranked by unique explained variance $\Delta R^2$ in predicting timescales.

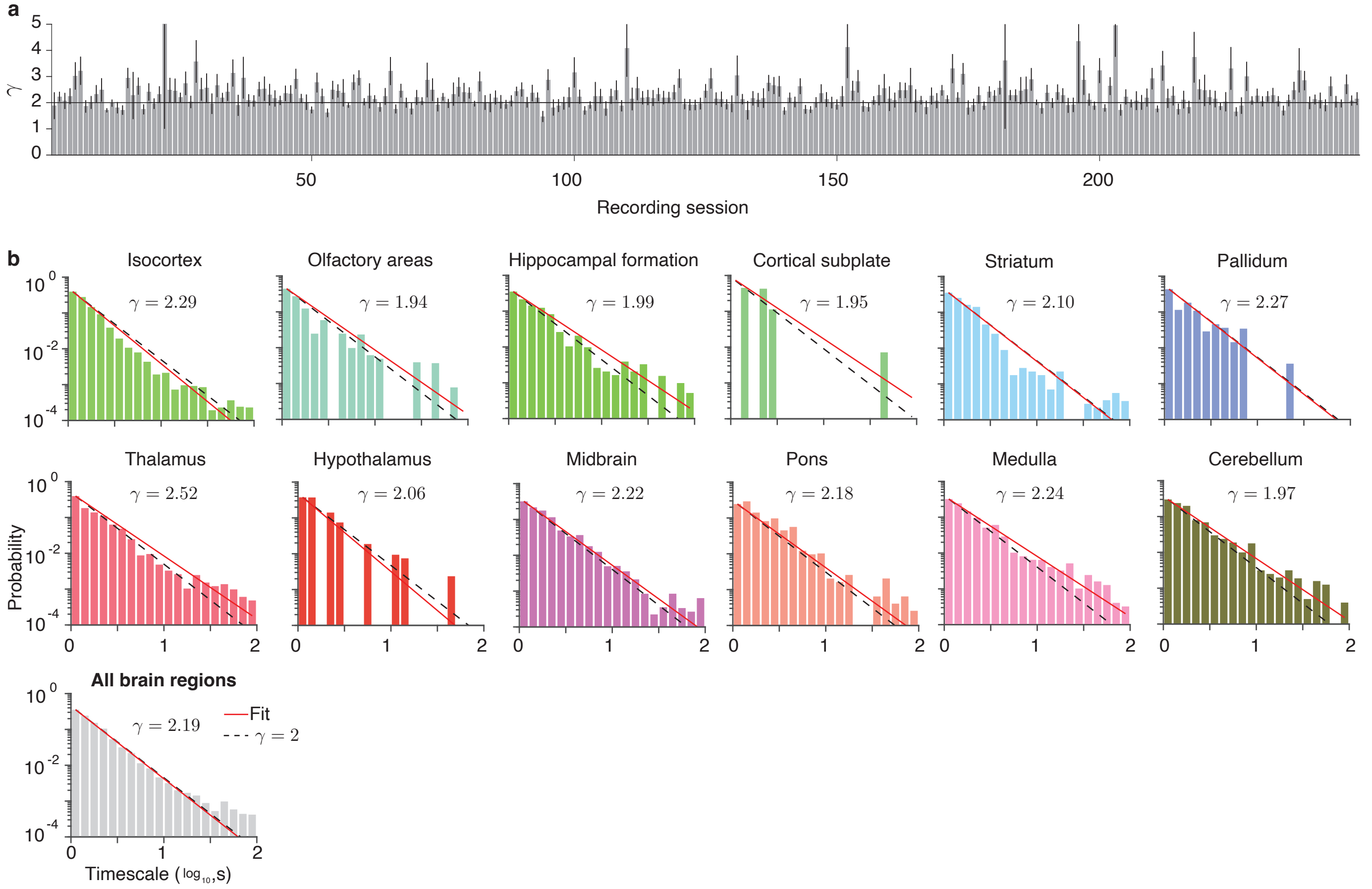


**Supplementary Figure 9. Additional tests of the power-law distribution of timescales across the brain.** **a**, The exponent $\gamma$ of the best-fitting power-law for timescale distributions was near 2 in most recording sessions, which had sufficient data for fitting (more than 40 neurons). Error bars represent 95% confidence intervals. The solid horizontal line indicates $\gamma = 2$. **b**, The distributions of timescales estimated from fitting a single-exponential function across neurons within each major brain structure (colored) and across neurons pooled from all regions (gray) are well described by a power law with an exponent $\gamma$ close to 2 (red line: best-fitting power law; dashed line: reference power law with exponent 2). All neurons were included in this analysis.

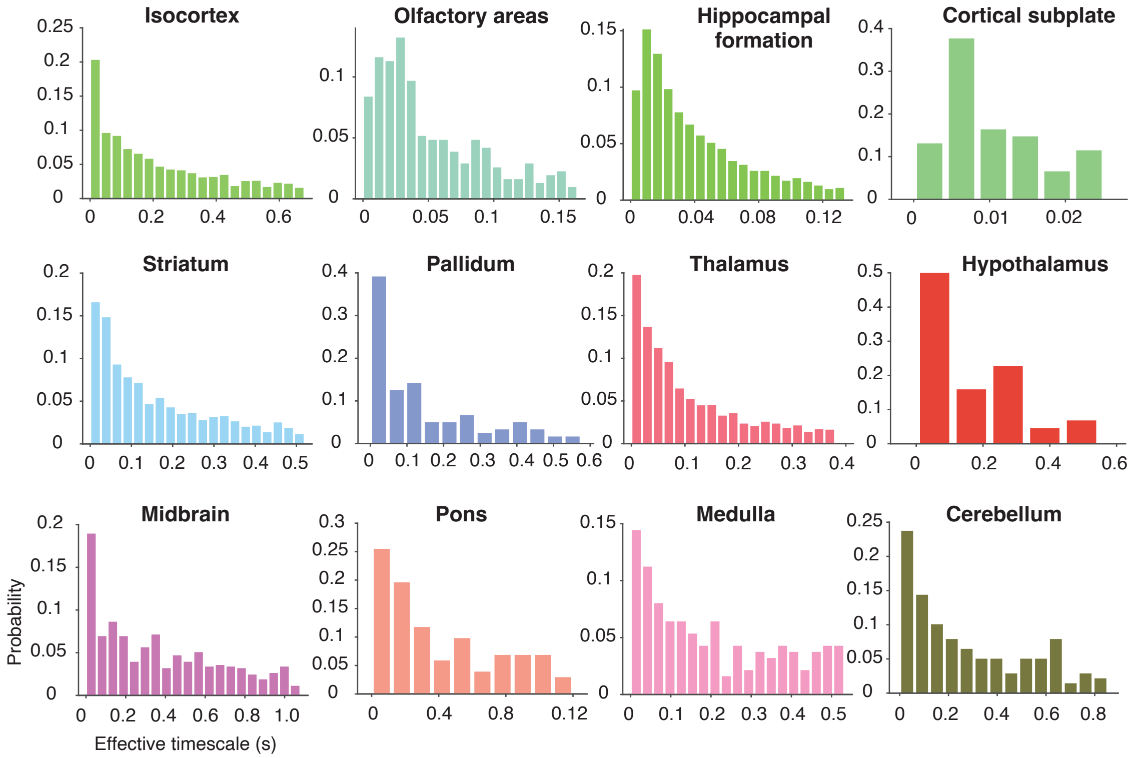


**Supplementary Figure 10. Distribution of fast timescales.** Distributions of fast timescales below the power-law cutoff, which do not follow a power law, for each of the 12 major brain structures.

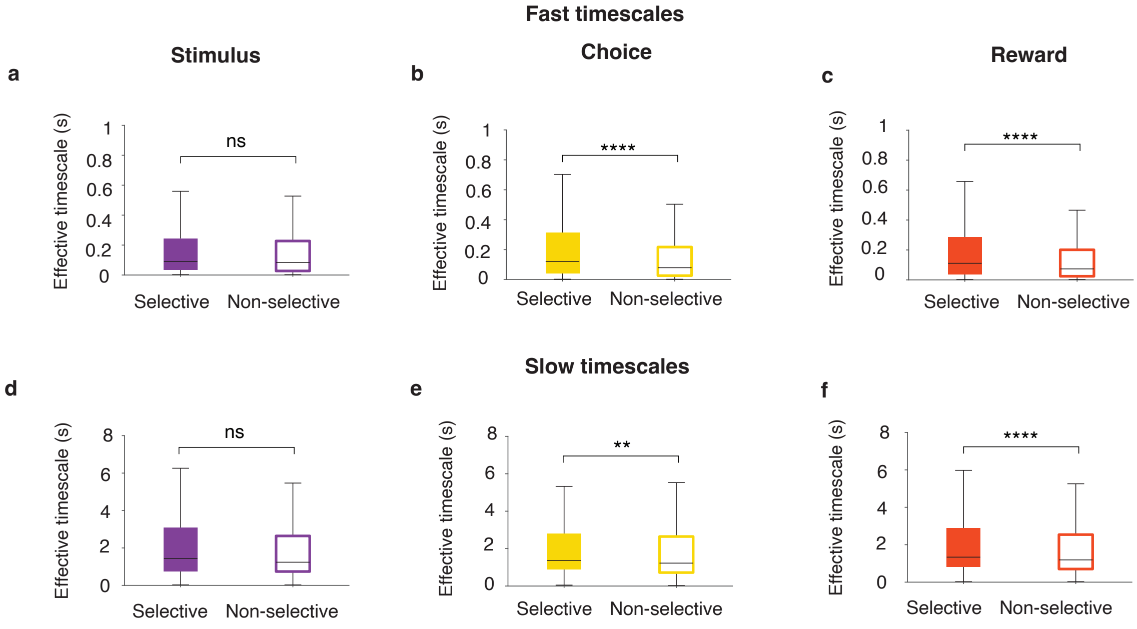


**Supplementary Figure 11. Relationships of fast and slow timescales with task-variable encoding. a**, Fast effective timescales did not differ significantly between neurons selective (filled bar) and non-selective (open bar) for the visual stimulus ($p = 0.09$, two-sided Wilcoxon rank sum test; $n = 490$ selective, $n = 6,480$ non-selective). In box plots (a-f), center lines indicate medians; boxes span the 25th to 75th percentiles; whiskers extend to the nearest of $1.5\times$ the interquartile range or the most extreme data point. **b**, Fast effective timescales were significantly longer in choice-selective neurons (filled bar) than in neurons non-selective for choice (open bar; $p < 10^{-10}$, two-sided Wilcoxon rank sum test; $n = 839$ selective, n = $6,131$ non-selective). **c**, Fast effective timescales were significantly longer in reward-selective neurons (filled bar) than in neurons non-selective for reward (open bar; $p < 10^{-10}$, two-sided Wilcoxon rank sum test; $n = 2,235$ selective, $n = 4,735$ non-selective). **d**, Slow effective timescales did not differ significantly between neurons selective (filled bar) and non-selective (open bar) for the visual stimulus ($p = 0.10$, two-sided Wilcoxon rank sum test; $n = 268$ selective, $n = 3,389$ non-selective). **e**, Slow effective timescales were significantly longer in choice-selective neurons (filled bar) than in neurons non-selective for choice (open bar; $p = 0.002$, two-sided Wilcoxon rank sum test; $n = 540$ selective, $n = 3,106$ non-selective). **f**, Slow effective timescales were significantly longer in reward-selective neurons (filled bar) than in neurons non-selective for reward (open bar; $p = 7.7 \cdot 10^{-6}$, two-sided Wilcoxon rank sum test; $n = 1,337$ selective, $n = 2,309$ non-selective).

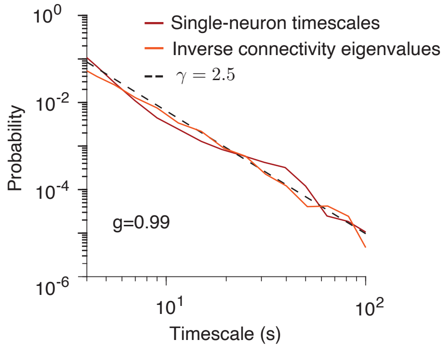


**Supplementary Figure 12. Comparison between the distributions of single-neuron timescales and inverse connectivity eigenvalues.** In linear networks, the distribution of single-neuron timescales, estimated by fitting their autocorrelation with an exponential function (red), closely matches the distribution of the network timescales given by the inverse eigenvalues of the connectivity matrix (orange). Simulations are shown for a linear network with connection weights sampled from a zero-mean normal distribution, corresponding to a semi-circle distribution of eigenvalues ($\zeta = 0.5$). The synaptic gain parameter is $g = 0.99$. Dashed line: reference power law with $\gamma = 2.5$.

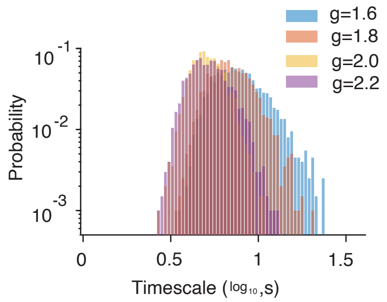


**Supplementary Figure 13. Distribution of timescales in nonlinear networks with Gaussian connectivity.** Distribution of timescales in nonlinear network models with connectivity weights sampled from a Gaussian distribution. For each value of $g$, we performed 1,000 simulations, following the procedures for timescale estimation described in the Methods.
